# EARLY-LIFE STRESS INDUCES SEXUALLY DIMORPHIC BRAIN AGING VIA MESOLIMBIC ALTERATIONS AND FRONTO-STRIATAL DYSFUNCTION

**DOI:** 10.64898/2026.10.09.757621

**Authors:** Amanda Cristina de Souza, Vinicius Schiavinatto Mariano, Isabele Cristina Gobo Tibiriça, Gabriel Boer Grigoletti-Lima, José Antonio Rocha Gontijo, Patrícia Aline Boer

## Abstract

Prenatal stress influences brain development; however, its long-term, region-specific effects on brain aging in males and females remain insufficiently characterized. This study examined the impact of unpredictable mild prenatal stress on behavior, cognition, and maternal care in C57BL/6J mice at middle (12 months) and old (18 months) ages.Stressed dams showed fragmented, unpredictable maternal behaviors, creating a stressful postnatal environment without impairing caregiving capacity. Both male and female offspring were born with lower birth weights and showed rapid postnatal growth, a trajectory associated with central insulin resistance and hippocampal signaling deficits. Sex-specific behavioral differences emerged with age. Middle-aged stressed males exhibited anhedonia, which progressed to hyperactivity, stereotypy, and persistent spatial memory deficits in old age. These outcomes corresponded with accelerated fronto-striatal decline, likely mediated by insulin resistance and increased rigidity of medial prefrontal cortex structures due to diminished local estrogenic activity. In contrast, middle-aged stressed females displayed hypoactivity and reduced anxiety, later developing anhedonia in old age while maintaining hippocampus-dependent memory. In females, this pattern supports a model in which a neuroestrogen - and growth factor-dependent feedback loop transiently preserves hippocampal plasticity but compromises dopaminergic function in reward circuits. This balance may deteriorate with reproductive senescence. Both sexes retained cognitive flexibility, indicating domain-specific rather than global cognitive impairment. These findings demonstrate that excessive prenatal glucocorticoid exposure and unpredictable postnatal care produce a combined hormonal and metabolic insult, resulting in sexually divergent trajectories of brain aging. The results underscore the necessity of incorporating sex as a biological variable in fetal programming research across the lifespan.

**HIGHLIGHTS:**

- Prenatal stress and fragmented care drive sexually dimorphic brain aging.
- Low birth weight and rapid growth associate with supposed central insulin resistance.
- Stressed males exhibit accelerated fronto-striatal decline and memory loss.
- Females show allostatic reallocation, transiently preserving hippocampal memory.

## 1 INTRODUCTION

Psychological stress constitutes a major global health issue, with an estimated 40 percent of adults worldwide reporting stress during a lot of the previous day in 2020 (Gallup, 2021). The prevalence of emotional stress has risen substantially, increasing from 26 percent in 2007 to nearly 38 percent in 2020 (Piao et al., 2024). The COVID-19 pandemic has further exacerbated rates of depression, anxiety, and other neuropsychiatric disorders (de Sousa Júnior et al., 2021; Wu et al., 2021; Yuan et al., 2022). Acute stress supports survival by mobilizing energy and enhancing focus through activation of the hypothalamic-pituitary-adrenal (HPA) axis (Ellis & Del Giudice, 2019; Gu et al., 2025; McEwen & Akil, 2020). In contrast, chronic stress leads to persistent HPA axis dysregulation and overloads the stress response system (Lei et al., 2025; Feng et al., 2025; Yang et al., 2025). During pregnancy, prolonged stress disrupts fetal brain development, resulting in lasting effects on stress regulation, metabolism, and emotional control. The Developmental Origins of Health and Disease (DOHaD) conceptposits that adverse prenatal conditions increase the risk of cardiovascular and metabolic diseases in adulthood (Barker, 1992; Barker, 1994; Barker, 1995; Hales & Barker, 1992). Recent research has extended this framework to encompass prenatal influences on brain development, linking gestational adversity to heightened risk of neuropsychiatric disorders (Fall, 2013; Sulyok et al., 2023; O’Donnell & Meaney, 2017). Excessive prenatal exposure to stress hormones leads to intrauterine growth restriction followed by rapid postnatal catch-up growth. This sequence is associated with metabolic dysfunction, termed “cerebral diabetes,” impaired synaptic plasticity, and hormone-mediated damage that accelerates brain aging (Wang et al., 2025; Grillo et al., 2019) during periods of intense cellular proliferation, migration, epigenetic plasticity, and fetal programming (Stevenson et al., 2020; Mirzakhani, 2024). Chronic gestational stress elevates circulating glucocorticoids beyond the protective capacity of placental 11β-hydroxysteroid dehydrogenase type 2 (11β-HSD2) (Mori & Shimosawa, 2025; Gotlieb et al., 2022; Eberle et al., 2021; Jensen Peña et al., 2012). Fetal glucocorticoid exposure disrupts neurogenesis and synaptogenesis and induces persistent epigenetic modifications (Kirkpatrick et al., 2025; Wu et al., 2025; Fowden et al., 2016). Prenatal stress disrupts the cyclical predictability of the neonatal environment, impeding the maturation of biological clocks and monoaminergic circuits and establishing a continuum of behavioral programming (Howland et al., 2021; Glynn & Baram, 2019). These effects affect neural circuits involved in emotional, cognitive, and motivational processes, particularly in the hippocampus, amygdala, prefrontal cortex (PFC), and mesolimbic dopaminergic system (Jagtap et al., 2023; Kim & Kim, 2023; Zoubovsky et al., 2022; Lautarescu et al., 2020). Male and female offspring respond differently to maternal stress, with male fetuses showing a reduced capacity to upregulate 11β-HSD2, resulting in greater glucocorticoid exposure in the male fetal brain (Bale, 2016; Bale & Epperson, 2015). Although prenatal stress induces enduring neurobiological changes, significant gaps remain in understanding how these effects manifest across the lifespan and differ by sex, particularly during midlife and old age, which are characterized by major endocrine changes (Bale, 2016; Bale & Epperson, 2015; McEwen & Morrison, 2013).

Three critical gaps require further investigation to delineate the temporal sequence of circuit-specific vulnerability: first, longitudinal studies assessing the effects of prenatal stress across both midlife and old age; second, clarification of sex-specific trajectories of brain aging across the lifespan; and third, integrated behavioral phenotyping—including reward processing, anxiety, spatial memory, cognitive flexibility, social behavior, and maternal care—within a single prenatal stress model. The present study addresses these gaps by characterizing the behavioral and cognitive outcomes of prenatal unpredictable chronic mild stress (UCMS) in male and female C57BL/6J mice at 12 months (midlife) and 18 months (old age). A comprehensive behavioral battery will assess reward processing, locomotor activity, anxiety and depression-like behavior, recognition memory, spatial learning, cognitive flexibility, and social behavior. We will also analyze maternal care fragmentation to provide an integrated perspective on the programming continuum shaping offspring neurodevelopmental trajectories. It is hypothesized that prenatal stress induces sex-specific trajectories of behavioral and cognitive aging. Specifically, male rodents are predicted to exhibit accelerated fronto-striatal decline, driven by impaired central metabolic disorders. In contrast, female rodents are predicted to demonstrate a compensatory reallocation of neural resources, with temporary preservation of hippocampal function.

## 2 MATERIALS AND METHODS

### 2.1. Ethical Statement and Animal Housing

All experimental procedures were conducted in accordance with the guidelines of the Brazilian National Council for the Control of Animal Experimentation (CONCEA) and were approved by the Ethics Committee for Animal Research at the State University of Campinas (CEUA: 5774-1/2021). Mice were housed in standard polypropylene cages (40 × 20 × 15 cm) containing wood shaving bedding and maintained under controlled environmental conditions (21 ± 2 °C; 12:12 h light–dark cycle, lights on at 06:00 h), with *ad libitum* access to water and standard laboratory chow (Nuvital®, Colombo, PR, Brazil). Measures were implemented to minimize animal suffering and to reduce the number of animals used.

### 2.2 Animals and Breeding

Male (n = 20) and female (n = 40) C57BL/6J mice were obtained from the Multidisciplinary Center for Biological Research (CEMIB) at the State University of Campinas. At six weeks of age, females were paired with males at a ratio of two females per male in individual breeding cages. Pregnancy was confirmed by the presence of a vaginal plug, which was designated as gestational day 0 (GD0). After pregnancy confirmation, males were removed, and females were housed individually.

### 2.3. Unpredictable Chronic Mild Stress (UCMS) Protocol

Pregnant dams were randomly assigned to either the control (C: n = 20) or stress (S: n = 20) groups. The stress group was subjected to an unpredictable chronic mild stress protocol from gestational day 6 (GD6) to gestational day 21 (GD21), as adapted from Guan et al. (2021) and Benoit et al. (2015). This protocol consisted of the daily application of a single stressor, rotated in a semi-random sequence to prevent habituation and maintain unpredictability. The stressors applied were as follows:

| Stressor | Duration |
| --- | --- |
| Lights on during the dark phase | Overnight (per 12 h) |
| Empty bedding cage (no bedding material) | 3 h |
| Cage tilted at 45° | 7 h |
| Food restriction | 24 h |
| Cage shaking (manual agitation) | 30 min |
| Water restriction | 24 h |
| Wet cage (200 mL water added to bedding) | 8 h |
| White noise (1500 Hz, 92 dB) | 1 h |
| Physical restraint (50 mL conical tube) | 4 h |
| Forced swim (31 °C water) | 1 h |

Each stressor was applied once daily, individually, and the sequence was repeated in a rotating order until birth. The control group was maintained under standard housing conditions without intervention, except for routine cage changes conducted at the same time of day as stressor application in the stress group.

The protocol includes metabolic stressors, specifically limiting food and water for 24 hours, which might complicate the effects of psychological stress and short-term nutritional deprivation. This approach matches established methods for unpredictable chronic mild stress (UCMS) in prenatal stress studies (48, 49). However, the possible impact of nutrition on the results cannot be completely ruled out and is discussed in the Limitations section.

### 2.4. Biometric Parameters

#### 2.4.1. Maternal Parameters

We recorded mothers’ body weight from GD0 to postnatal day 21 (PND21), and measured food intake at PND7 (C: n = 10; S: n = 14). The stillbirth rate (C: n = 14; S: n = 14) and the number of pups born per litter (C: n = 20; S: n = 20) were recorded at birth.

#### 2.4.2. Offspring Parameters

We weighed offspring of both sexes at PND0 (birth), PND7, PND14, and PND21 to assess the prenatal stress protocol and track postnatal growth. At PND1, we reduced litters to a maximum of eight pups per mother to ensure equal access to food.

### 2.5. Maternal Behavior Assessment

We adapted the protocol from Molet et al. (2016) and Zoubovsky et al. (2020). We studied maternal behavior in a group of mothers (C: n = 10; S: n = 10) from PND2 to PND9. Observations took place during the light period in two daily 30-minute sessions: one in the morning (09:00–10:30) and one in the afternoon (14:00–15:30). Before the behavior tests, mothers acclimated to the experimental room for 30 minutes. No cage changes happened during either the getting-used-to or testing times. During each observation session, the duration (in seconds) of the following maternal behavioral parameters was recorded:

1.Licking/Grooming (LG): Time spent licking and grooming pups.
2.Arched-back Nursing (ABN): Nursing in an active arched-back posture.
3. Passive Nursing (PN): Nursing in a passive, flat-backed position.
4. Off-Nest (ON): Time spent away from the nest.
5. Nest Building (NB): Time spent constructing or rearranging the nest.
6. Self-Grooming (SG): Time spent grooming herself.

Four independent observers recorded the duration of each behavior using the smartphone app Countee (Slanzi & Fernand, 2024). Observers checked agreement by calculating the intraclass correlation coefficient (ICC) on some recordings. For analysis, we compared total time spent on each behavior between groups.

#### 2.5.1. Behavioral Fragmentation Analysis

We measured behavioral fragmentation as a sign of unpredictability and timing problems in maternal care, following Molet et al. (2016) and Baram et al. (2012). For each behavior, fragmentation was calculated by dividing the total number of behavior episodes by the average length of each episode, giving a fragmentation index (FI):

FI = Total number of bouts / Mean bout duration (s)

Higher FI values indicate greater fragmentation, as evidenced by more frequent, shorter episodes that reflect a less predictable, more disorganized caregiving style. We measured fragmentation for licking/grooming, nest building, off-nest, passive nursing, and self-grooming behaviors.

#### 2.5.2. Pup Retrieval Test

We adapted the pup retrieval test from Wang and Storm (2011). On PND3, we briefly removed the mother from the home cage, then randomly selected three pups and placed them on the side of the cage opposite the nest: two in the corners and one in the middle. We then returned the mother to the cage between the nest and the scattered pups. We recorded the time it took to pick up the first pup immediately after returning the mother. Timing for the first pup ended when the mother put it in the nest, and timing for the second pup began. We repeated this for all three pups. The maximum allowed time to gather all pups was 600 seconds. For analysis, we compared the time to get the first pup and the total time to get all three pups between groups.

### 2.6. Behavioral Testing of Offspring

Following weaning at postnatal day 21 (PND21), offspring were housed in same-sex groups of three to five per cage under standard laboratory conditions until behavioral testing. Behavioral assessments were conducted at 12 months (middle age) and 18 months (old age), with one offspring per mother (male or female) assigned to each group. To reduce potential carryover effects between tests, offspring were allocated to one of two distinct groups:

- Cohort 1 (Barnes Maze): Animals in this group completed only the 10-day Barnes maze protocol.
- Cohort 2 (Behavioral Battery): Animals in this group underwent a sequential battery of tests, including the sucrose preference test, open field activity monitoring, novel object recognition test, elevated plus maze, and social interaction test. All behavioral assessments were conducted during the light phase (08:00–14:00) in a dedicated testing room maintained at 22 ± 1 °C, with background noise below 50 dB and dim lighting (30–50 lux, unless otherwise specified). Equipment was sanitized with 70% ethanol between trials to eliminate olfactory cues. Testers were blinded to group assignments whenever possible. Animals were acclimated to the testing room for at least 30 minutes before each session.

#### 2.6.1. Sucrose Preference Test

The sucrose preference test assessed anhedonia-like behavior, a principal indicator of depression-like traits in rodents (Markov, 2022; Berrio et al., 2024). Female (12 months: Cn = 17, S n = 17; 18 months: Cn = 14, S n = 16) and male (12 months: Cn = 10, S n = 11; 18 months: Cn = 15, S n = 12) animals were individually housed during testing.

The protocol consisted of three consecutive days:

- Day 1 (Adaptation): We provided two identical bottles filled with water to allow habituation to the two-bottle paradigm.
- Day 2 (Test 1): We filled one bottle with water and the other with a 1% (w/v) sucrose solution, then positioned the bottles randomly.
- Day 3 (Test 2): We repeated the procedure, swapping bottle positions to control for side preference.

In accordance with current guidelines for sucrose preference testing in rodents (Berrio et al., 2024), animals were not deprived of food or water before testing. We quantified fluid intake by weighing the bottles every 24 hours. Sucrose preference was calculated as:

Sucrose Preference (%) = [Sucrose intake / (Sucrose intake + Water intake)] × 100

Total fluid intake (sucrose plus water) was also recorded to account for potential differences in overall consumption between groups.

#### 2.6.2. Activity Monitor (Open Field) Test

Movement and exploration were measured using an open field box (45 × 45 × 45 cm; model ENV-515; Med Associates Inc., St. Albans, VT, USA) with two 16-beam infrared sensors that automatically tracked side-to-side and up-and-down movements. Animals were placed in the center and allowed to explore freely for 5 minutes in dim light (30 lux) inside a quiet room.

The following parameters were recorded and analyzed: Distance traveled (cm); Average speed (cm/s); Time spent in the safe zone (peripheral area); Ambulatory movements (horizontal beam breaks); Unstable or stereotyped movements; Rotational movements; Number of jumps; Number of rearing events (standing upright); Rest time (s).

#### 2.6.3. Novel Object Recognition (NOR) Test

The NOR test assessed short- and long-term recognition memory, based on rodents’ innate preference for novel stimuli (Antunes & Biala, 2012). Females (12 months: C n = 14, S n = 13; 18 months: C n = 13, S n = 15) and males (12 months: C n = 12, S n = 11; 18 months: C n = 9, S n = 11) were tested.

The protocol consisted of three phases:

• Day 1 (Habituation): Animals were placed in the empty apparatus (40 × 40 × 40 cm) for 5 minutes to habituate to the environment.

• Day 2, Morning (Familiarization): Two identical objects (Object A1 and A2) were placed in opposite corners of the apparatus. Animals explored for 5 minutes.

• Day 2, Afternoon (Short-term memory test – 3 h delay): One familiar object was replaced with a novel object (Object B). Animals explored for 5 minutes.

• Day 3 (Long-term memory test – 24 h delay): The familiar object was again replaced with a different novel object (Object C). Animals explored for 5 minutes.

Object positions and identities were counterbalanced across animals to control for object and location preferences. Exploration was defined as directing the nose toward the object within ≤2 cm. The discrimination index (DI) was calculated as:

DI = (Time exploring novel object − Time exploring familiar object) / Total exploration time × 100 2.6.4. Elevated Plus-Maze (EPM) Test

Anxiety-like behavior was measured using the elevated plus maze, a well-established test of approach and avoidance behavior in rodents (Walf & Frye, 2007). The setup had two open arms (30 × 5 cm) and two closed arms (30 × 5 × 15 cm), connected by a central square platform (5 × 5 cm), and raised 50 cm above the floor. Females (12 months: C n = 11, S n = 18; 18 months: C n = 14, S n = 16) and males (12 months: C n = 8, S n = 10; 18 months: C n = 14, S n = 11) were placed on the central platform facing an open arm and allowed to explore for 5 minutes under dim red light (5 lux). The following parameters were recorded: Time spent in open arms (s); Time spent in closed arms (s); Time spent in the central platform (s); Number of entries into open arms; Number of entries into closed arms; Total number of arm entries (open + closed); The percentage of time spent in open arms was calculated as: % Open Arm Time = [Time in open arms / (Time in open arms + Time in closed arms)] × 100 We used the total number of arm entries to measure general movement and rule out motor-related effects.

#### 2.6.5. Social Interaction Test

We tested social behavior using a three-chamber setup. Removable walls divided the box into three sections. Females (12 months: C n = 13, S n = 12; 18 months: C n = 15, S n = 12) and males (12 months: C n = 12, S n = 10; 18 months: C n = 15, S n = 12) were tested.

The protocol consisted of three phases:

- Phase 1 (Habituation): The test animal was placed in the apparatus for 10 minutes with empty wire cups in both side chambers.
- Phase 2 (Sociability): After a 45-second interval, the test animal was placed in the center compartment, with the partitions closed, for 3 minutes. An unfamiliar age- and sex-matched stimulus mouse (Stranger 1) was placed in a wire cup in one side chamber, while the other cup remained empty. Partitions were then opened, and the test animal was allowed to explore for 5 minutes.
- Phase 3 (Social Novel Preference): After another 45-second interval, the test animal was again placed in the center, with the partitions closed, for 3 minutes. The familiar mouse (Stranger 1) remained in its original position, and a novel, unfamiliar mouse (Stranger 2) was placed in the opposite wire cup. Partitions were opened, and the test animal was allowed to explore for 5 minutes. The following parameters were recorded:
- Time spent in each chamber
- Time spent sniffing/investigating each stimulus mouse
- Sociability index = Time with Stranger 1 / (Time with Stranger 1 + Time with empty cup)
- Social novelty index = Time with Stranger 2 / (Time with Stranger 1 + Time with Stranger 2)

#### 2.6.6. Barnes Maze Test

The Barnes maze tested spatial learning, memory, and mental flexibility. It had a round platform (90 cm wide) with 20 evenly spaced holes (5 cm wide) around the edge. We placed a black escape box (9 × 20 × 9 cm) under one hole (the target). We placed four different visual signs (square, circle, triangle, and cross) on the walls around the maze. We conducted testing under bright light (1200 lux) to encourage escape behavior.

Females (C: n = 14; S: n = 15) and males (C: n = 14; S: n = 15) aged 18 months were tested. The protocol consisted of the following phases:

Day 1 – Habituation: Animals explored the platform freely for 1 minute, then were gently guided to the escape box and left undisturbed for 2 minutes.

Days 2–6 – Learning the location: Each animal was placed in the center of the platform inside a black opaque cylinder for 15 seconds. After the cylinder was removed, the animal could explore for up to 180 seconds to find the escape hole. Once it found it, the animal stayed in the escape box for 30 seconds. Three trials were done each day with 15 minutes between trials. The time taken to find the escape hole was recorded for each trial.

Day 7 – Probe Trial (Memory Assessment): The escape box was removed. After 15 seconds in the center, the animal explored the platform for 90 seconds. The percentage of time spent in the target quadrant (the quadrant containing the former escape hole location) was recorded as an index of spatial memory retention.

Days 8–10 – Reversal Learning (Cognitive Flexibility): The escape box was relocated to the diametrically opposite hole (180° rotation). The same acquisition procedure was followed (three trials/day, 180 s maximum, 15-min inter-trial interval).

Day 11 – Probe Trial (Cognitive Flexibility Assessment): The escape box was removed, and a 90-second probe trial was conducted as on Day 7.

Search Strategy Classification - Search strategies were classified based on the trajectory pattern used to locate the escape hole, using an adaptation of the Barnes maze search-strategy framework of Rodríguez Peris et al. (2024), with the following operational categories:

- Direct strategy: The animal reached the target hole directly with ≤2 errors.
- Corrected strategy: The animal approached the target region, making 3–5 errors before locating the hole.
- Serial strategy: The animal searched holes sequentially around the perimeter.
- Random strategy: The animal exhibited no systematic search pattern.

We assigned a cognitive score (1–11) to each trial based on search-strategy efficiency, with higher scores indicating more efficient (direct) strategies.

Approach Index: We calculated this as the total distance traveled to reach the target area (a 15-cm radius around the escape hole) during each trial. Higher numbers mean less efficient navigation.

### 2.7. Statistical Analysis

Data are shown as mean ± standard deviation (SD). We checked whether the data were normally distributed using the Shapiro–Wilk test and whether the variances were equal using Levene’s test. Data that met these conditions were analyzed with Student’s *t*-test (for two groups), multiple *t*-tests with Holm–Šidák correction (for multiple comparisons), or two-way ANOVA (factors: group × age/sex) followed by Tukey’s post hoc test. We analyzed data that did not meet these conditions using the Mann–Whitney *U* test. Results were considered significant if *p* < 0.05. All analyses were done using GraphPad Prism software (version 10; GraphPad Software, San Diego, CA, USA).

## 3. RESULTS

### 3.1. Maternal Biometric Parameters

During pregnancy (GD0–GD20), body weight was similar between stressed (S) and control (C) mothers (Figure 1A). However, on the day they gave birth, S mothers weighed significantly less than C mothers (Figure 1A). From PND7 to PND21, body weight stayed similar between the two groups (Figure 1A). At PND7, S mothers ate significantly more food than C mothers (Figure 1B). Food intake did not differ significantly at other times. Stress before birth caused a significant rise in stillbirths in the S group compared to the C group (Figure 1C). In addition, the number of pups per litter was much lower in the S group than in the control group (Figure 1D).

**Figure 1.**
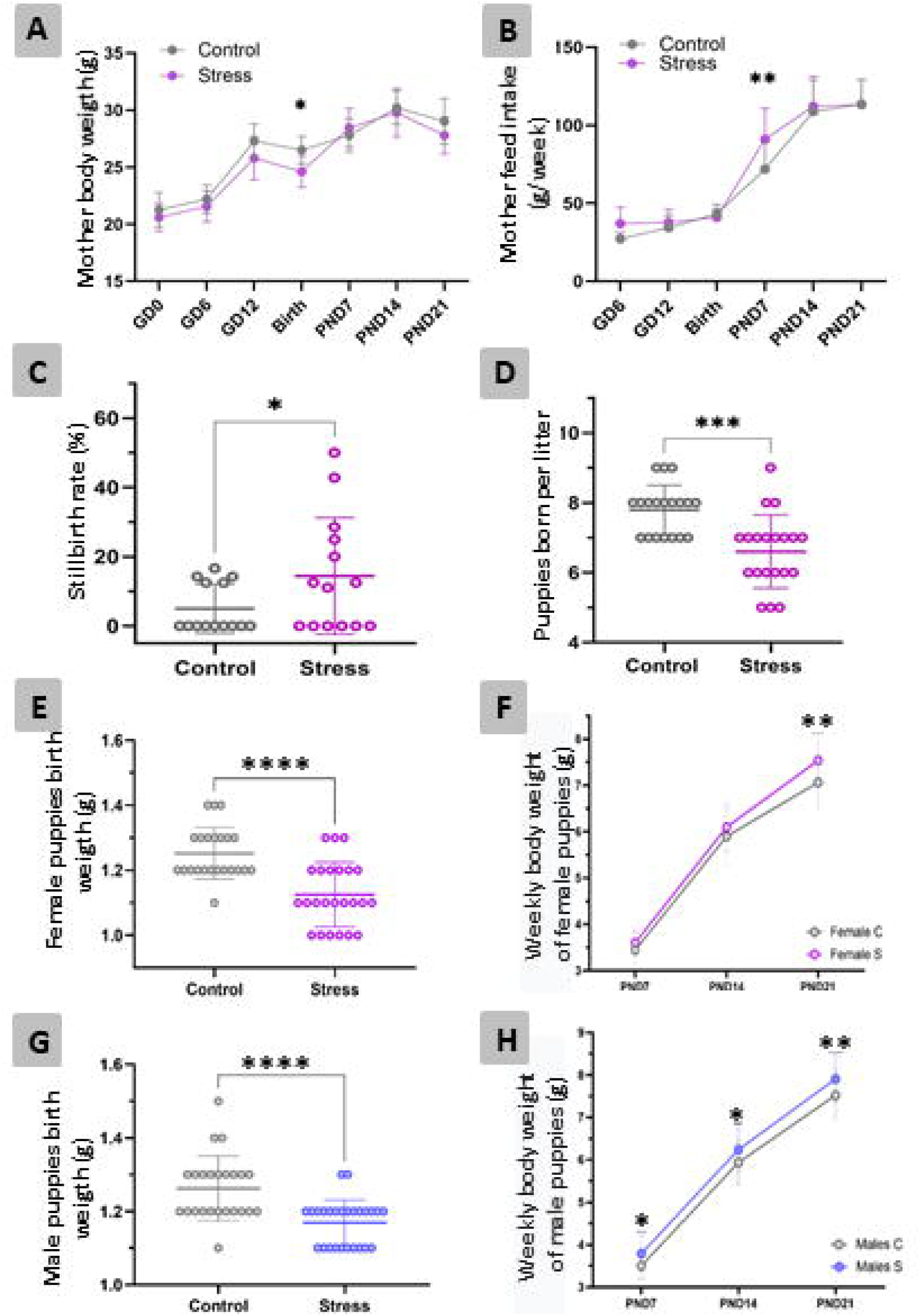
Effects of gestational stress on maternal and offspring weight and reproductive outcomes. Maternal body weight during GD 0 until PND 21 (A) and food intake of mothers during GD 6 until PND 21 (B). No differences were observed during gestation; however, the S group showed a significant reduction in body weight at parturition. (B) Food intake measured at PND7. (C) Stillbirth rate (%) in control and prenatal stress groups. (D) Number of pups born per litter in control and prenatal stress groups. Prenatal stress increased the stillbirth rate and reduced litter size compared with controls. Offspring body weight at PND0 in female (E) and male (G) offspring from control and stress groups. (B, D) Body weight at birth (E and G) and during development at PND7, PND14, and PND21 in female (F) and male (H) offspring. Data are presented as mean ± SD. *p < 0.05, **p < 0.01, **p < 0.001, ****p < 0.0001.

### 3.2. Offspring Biometric Parameters

Female offspring in the S group exhibited significantly lower body weight at birth (PND0) compared to the C group (Figure 1E). Subsequently, body weight in the S group increased steadily, with S females weighing significantly more than C females at PND7, PND14, and PND21 (Figure 1F). These findings indicate accelerated postnatal growth in the S group. A comparable trend was observed in male offspring. S males demonstrated significantly lower birth weight at PND0 compared to C males (Figure 1G). However, at PND7, PND14, and PND21, S males exhibited significantly higher body weight than controls (Figure 1H).

### 3.3. Maternal Behavior

#### 3.3.1. Duration of Maternal Behaviors

The Student’s *t*-test revealed significant differences in the duration of maternal behaviors across groups (Figure 2). S mothers spent much less time building nests (Figure 2A) and licking or grooming their pups (Figure 2D) than C mothers. On the other hand, S mothers spent much more time grooming themselves (Figure 2C), and quietly nursing (Figure 2E) compared to C mothers. Groups did not differ in time spent eating or drinking (Figure 2B. These findings show that the main motherly duties remained the same, even though some caregiving behaviors changed.

**Figure 2.**
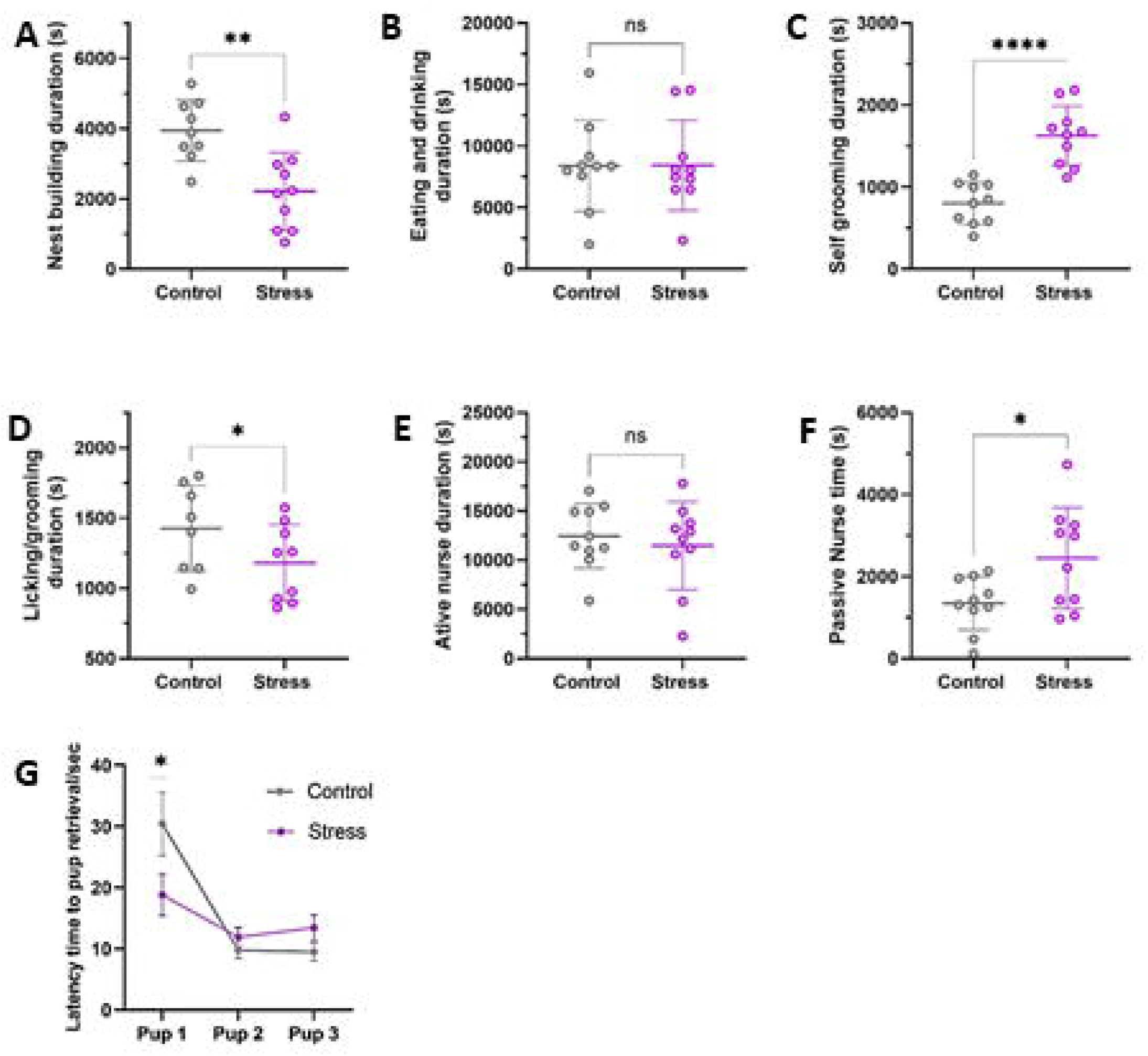
Effects of gestational stress on maternal behavior. Nest building duration (A), eating and drinking duration (B), self-grooming duration (C), licking/grooming duration (D), active nursing duration (E), and passive nursing time (F) were evaluated in control and prenatal stress dams. (G) Latency time to mother retrieval of offspring. Data are presented as mean ± SD *p < 0.05, **p < 0.01, ****p < 0.0001; ns = not significant.

#### 3.3.2 Pup Retrieval Assessment

On postnatal day 3, S mothers required significantly less time to retrieve the first pup than C mothers (Figure 2G). No significant differences were observed in the total time to return all three pups to the nest, or in the time required to retrieve the second or third pup (Figure 2G).

#### 3.3.3. Behavioral Fragmentation and Predictability

Given how predictable behavior was, S dams showed less predictable overall behavior than C dams (Figure 3K, L), which fits with a more disorganized, fragmented caregiving style. Looking at how broken-up behaviors were, there were clear differences between groups (Figure 3). S dams showed more broken-up behavior, as evidenced by more frequent, shorter episodes, in off-nest activity (Figure 3E, F), passive nursing (Figure 3I, J, and self-grooming (Figure 3A, B), than C dams. On the other hand, broken-up nest building was much less in S dams than in controls (Figure 3G, H). Broken-up licking/grooming did not differ clearly between groups (Figure 3C, D). Counts of each behavior showed no clear differences between groups in the total number of licking/grooming, nest building, off-nest, passive nursing, or self-grooming episodes. This suggests that differences in broken-up behavior stem from changes in episode length, not episode number.

**Figure 3.**
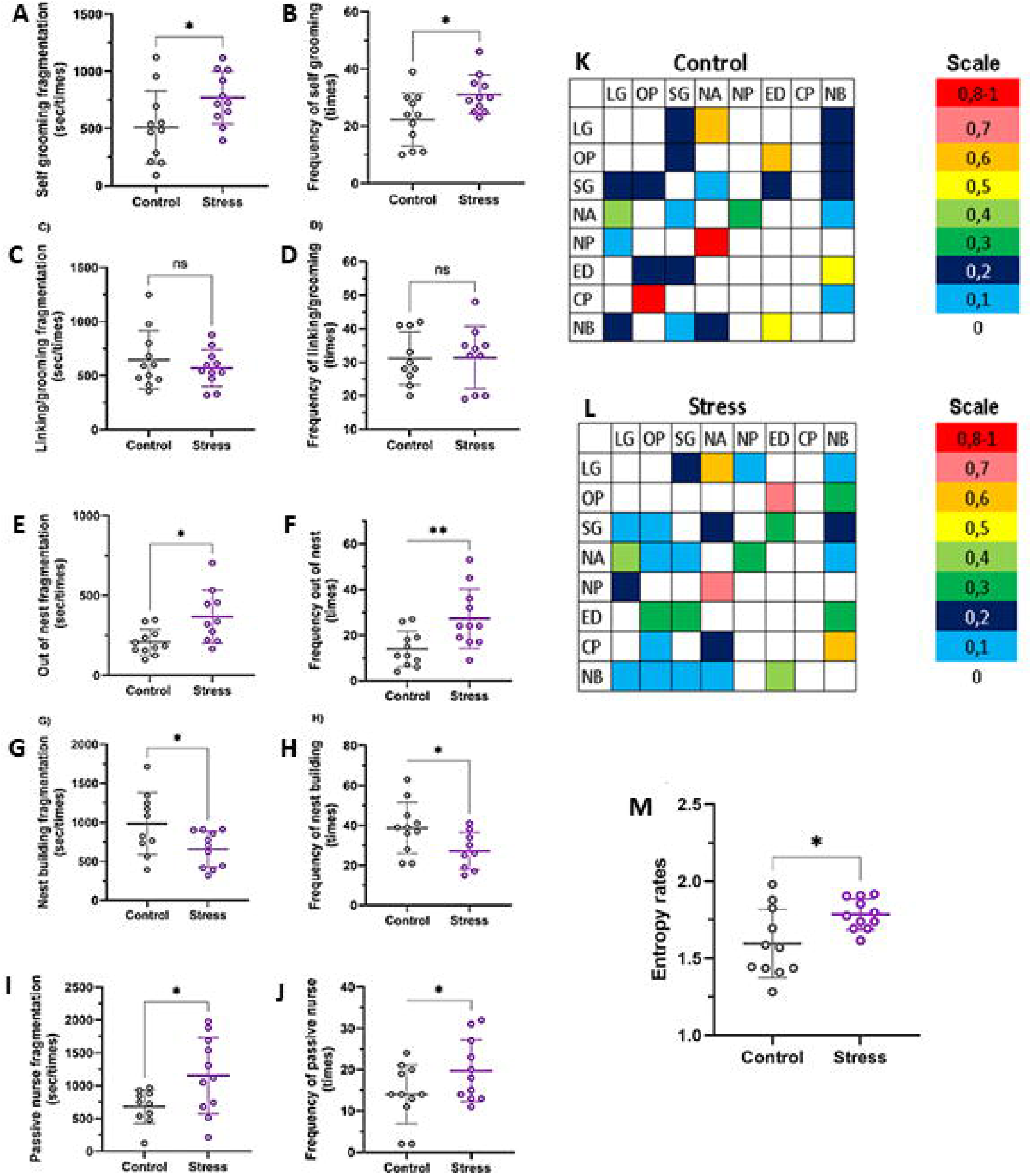
Maternal behavior and behavioral sequence organization and entropy rates. Self-grooming fragmentation (A) and frequency of self-grooming episodes (B), linking/grooming fragmentation (C) and frequency (D), out-of-nest fragmentation (E) and frequency of leaving the nest (F), nest-building fragmentation (G) and frequency of nest-building episodes (H), passive nursing fragmentation (I), and frequency of passive nursing behavior (J) were evaluated in control and stress groups. Transition matrices representing behavioral sequence probabilities in control (K) and stress dams (L) demonstrate alterations in the organization and dynamics of maternal behavioral patterns. Colors indicate transition probabilities between behavioral categories, ranging from low (blue) to high (red) frequencies. Entropy rates analysis (M) revealed significantly increased entropy in stressed dams, indicating greater behavioral unpredictability, reduced stereotypy, and impaired organization of maternal behavioral sequences. Data are presented as mean ± SD p < 0.05.* Data are presented as mean ± SD. *p < 0.05, p < 0.01, ns = not significant.

### 3.4. Sucrose Preference Test

Prenatal stress affected sucrose preference depending on sex and age (Figure 4). In females, there was no clear difference in sucrose preference between C and S groups at 12 months (Figure 4). At 18 months, S females showed a noticeably lower sucrose preference than C females (Figure 4). At 12 months, S males had a significantly lower sucrose preference than C males (Figure 4). At 18 months, no clear difference in sucrose preference was found between male groups (Figure 4).

**Figure 4.**
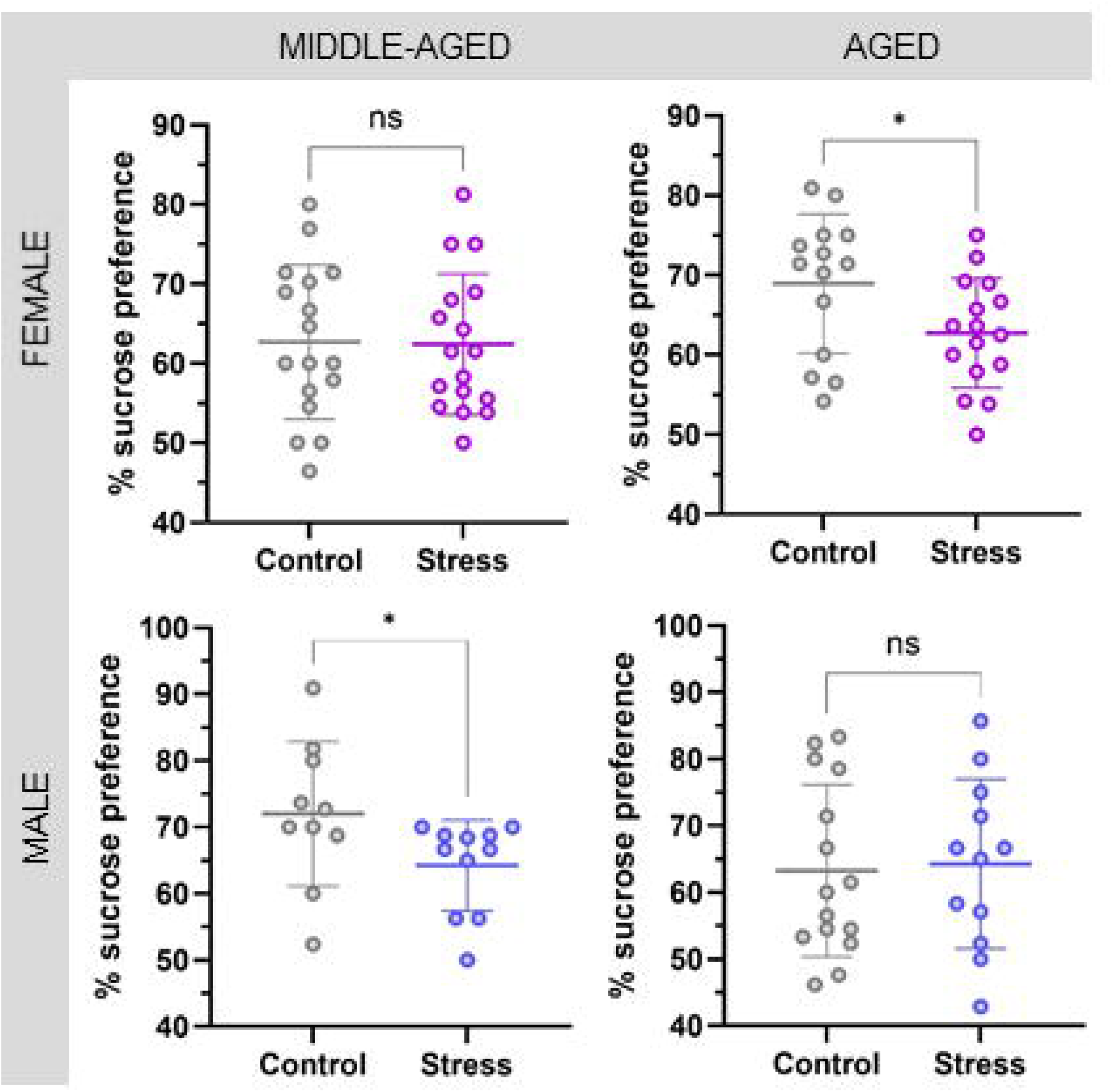
Sucrose preference test in male and female offspring at 12 and 18 months of age. Data are presented as mean± SD. *p<0.05.

### 3.5. Activity Monitor (Open Field) Test

The activity monitor test showed sex- and age-based differences in movement and exploration (Table I). At 12 months, S females moved a shorter distance, had a slower average speed, spent less time in the active area, made fewer walking movements, and rested more than C females, showing less overall activity. At 18 months, no clear differences in movement or exploration were found between C and S females. At 12 months, S males jumped less than C males, indicating less vertical exploration; no other major differences were observed between male groups at this age. At 18 months, S males moved a longer distance, had a faster average speed, spent more time in the active area, made more walking movements, and rested less than C males. In addition, S males showed more exploratory behaviors such as standing upright longer and jumping more often.

**Table I.**
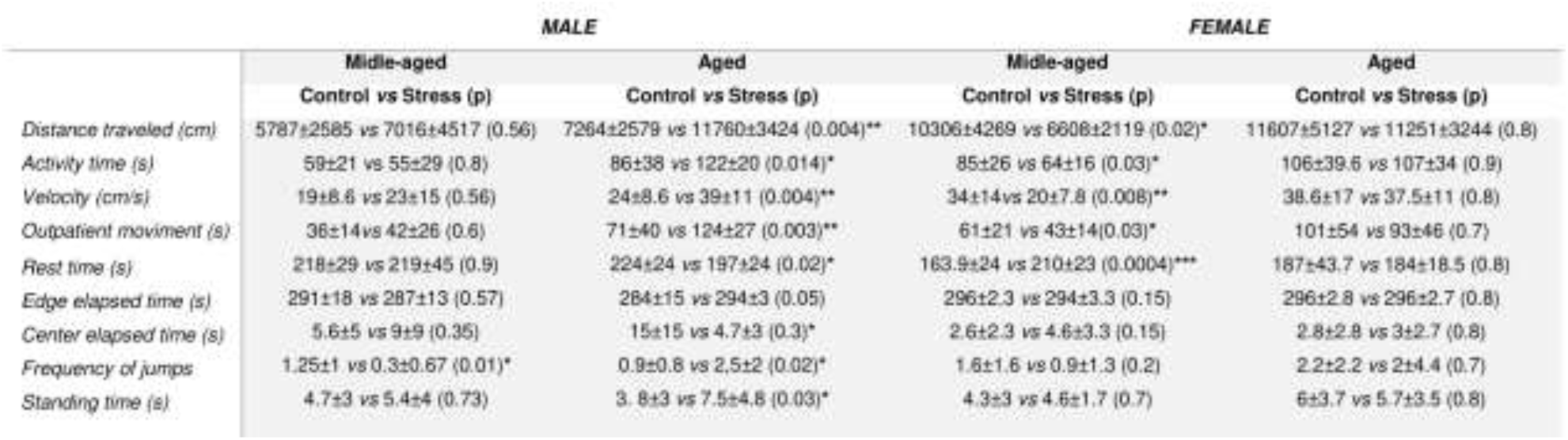
Activity monitor (open field) test results in offspring at 12 and 18 months of age. Data are presented as mean ± SD. Effect sizes (Cohen’s d) are reported for key comparisons to facilitate interpretation independent of sample size, particularly for underpowered groups. *p < 0.05, **p < 0.01, ***p < 0.001.

|  | MALE |  | FEMALE |  |
| --- | --- | --- | --- | --- |
|  | Middle-aged | Aged | Middle-aged | Aged |
|  | Control vs Stress (p) | Control vs Stress (p) | Control vs Stress (p) | Control vs Stress (p) |
| Distance traveled (cm) | 5787±2585 vs 7016±4517 (0.56) | 7264±2579 vs 11760±3424 (0.004)** | 10306±4289 vs 6608±2119 (0.02)* | 11607±5127 vs 11251±3244 (0.8) |
| Activity time (s) | 59±21 vs 55±29 (0.8) | 86±38 vs 122±20 (0.014)* | 85±26 vs 64±16 (0.03)* | 106±39.6 vs 107±34 (0.9) |
| Velocity (cm/s) | 19±8.6 vs 23±15 (0.56) | 24±8.6 vs 39±11 (0.004)** | 34±14 vs 20±7.8 (0.008)** | 38.6±17 vs 37.5±11 (0.8) |
| Outpatient movement (s) | 36±14 vs 42±26 (0.6) | 71±40 vs 124±27 (0.003)** | 61±21 vs 43±14 (0.03)* | 101±54 vs 93±46 (0.7) |
| Rest time (s) | 218±29 vs 219±45 (0.9) | 224±24 vs 197±24 (0.02)* | 163.9±24 vs 210±23 (0.0004)*** | 187±43.7 vs 184±18.5 (0.8) |
| Edge elapsed time (s) | 291±18 vs 287±13 (0.57) | 284±15 vs 294±3 (0.05) | 296±2.3 vs 294±3.3 (0.15) | 296±2.8 vs 296±2.7 (0.8) |
| Center elapsed time (s) | 5.6±5 vs 9±9 (0.35) | 15±15 vs 4.7±3 (0.3)* | 2.6±2.3 vs 4.6±3.3 (0.15) | 2.8±2.8 vs 3±2.7 (0.8) |
| Frequency of jumps | 1.25±1 vs 0.3±0.67 (0.01)* | 0.9±0.8 vs 2.5±2 (0.02)* | 1.6±1.6 vs 0.9±1.3 (0.2) | 2.2±2.2 vs 2±4.4 (0.7) |
| Standing time (s) | 4.7±3 vs 5.4±4 (0.73) | 3.8±3 vs 7.5±4.8 (0.03)* | 4.3±3 vs 4.6±1.7 (0.7) | 6±3.7 vs 5.7±3.5 (0.8) |

### 3.6. Elevated Plus-Maze Test

The elevated plus maze test showed changes in anxiety-like behavior depending on sex and age (Table II). At 12 months, S females spent more time in the open arms and less time in the closed arms than C females, showing less anxiety-like behavior. There were no significant differences in time spent on the central platform. For 12-month-old males, we found no significant differences between the C and S groups in any measure. At 18 months, S males spent more time in the open arms and less time in the closed arms than C males, also showing less anxiety-like behavior. No significant differences were seen between C and S females at 18 months in any measure.

**Table II.**
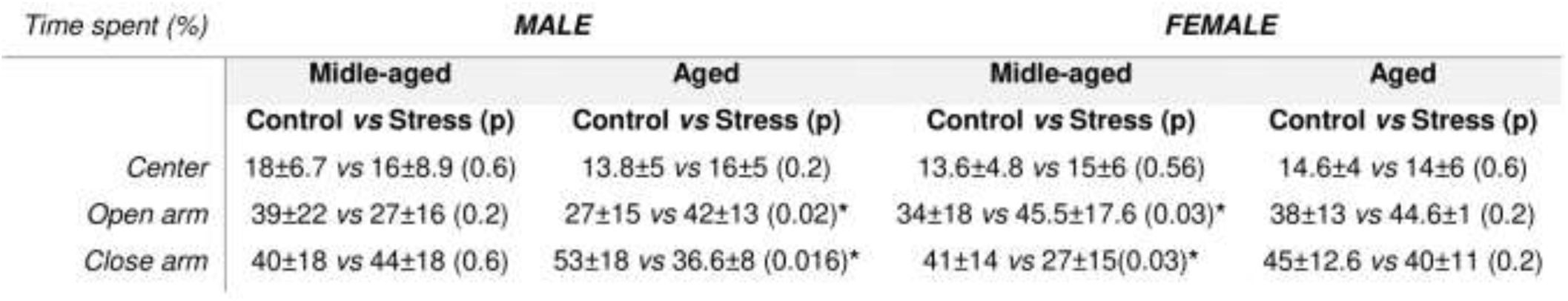
Elevated plus maze performance in offspring at 12 and 18 months of age. Data are presented as mean ± SD. Effect sizes (Cohen’s d) are reported for key comparisons to facilitate interpretation independent of sample size, particularly for null findings where statistical power is limited. *p < 0.05.

**Table III.** Approach index (distance to target zone) in the Barnes maze for aged male offspring (18 months). Data are presented as mean ± SD. *p < 0.05.

|  | MALE | FEMALE |
| --- | --- | --- |
|  | Control vs Stress (p) | Control vs Stress (p) |
| Day 1 | 544737±76718 vs 589899±94363 (0.2) | 593621±88602 vs 593805±110400 (0.99) |
| Day 2 | 432948±140758 vs 550395±140001 (0.04)* | 374732±147067 vs 453027±147459 (0.09) |
| Day 3 | 270122±177965 vs 431199±147552 (0.02)* | 294874±227853 vs 341120±159833 (0.5) |
| Day 4 | 225233±146511 vs 276851±107222 (0.3) | 217512±186414 vs 306468±190588 (0.2) |
| Day 5 | 153292±109621 vs 239936±132713 (0.04)* | 199836±205953 vs 210732±154710 (0.8) |
| Day 7 | 222871±138929 vs 195281±104707 (0.6) | 209417±207916 vs 200725±148235 (0.9) |
| Day 8 | 173328±104439 vs 223123±128576 (0.3) | 178541±175522 vs 164545±126526 (0.8) |
| Day 9 | 96266±65590 vs 190120±132271 (0.03)* | 1287034±138350 vs 127436±74692 (0.98) |

### 3.7. Novel Object Recognition Test

The novel object recognition test checked both short-term (3 hours) and long-term (24 hours) memory (Figure 5). The discrimination index did not differ significantly between the C and S groups for either sex at 12 or 18 months in the short-term memory test (Figure 5A, C, E and G), indicating that short-term recognition memory remained unchanged across groups. The long-term memory test results at 24 hours are as follows:

- At 12 months, S females showed much better object recognition than C females (p < 0.05; Figure 5D), showing improved long-term memory.
- At 12 months, S males had much more trouble recognizing objects than C males (p < 0.05; Figure 5H), showing weaker long-term memory.
- At 18 months, S females still did better than C females in recognizing objects (*p* < 0.001; Figure 5K). In contrast, C females at this age did not prefer the new object over the familiar one, suggesting that their object recognition declined with age.

**Figure 5.**
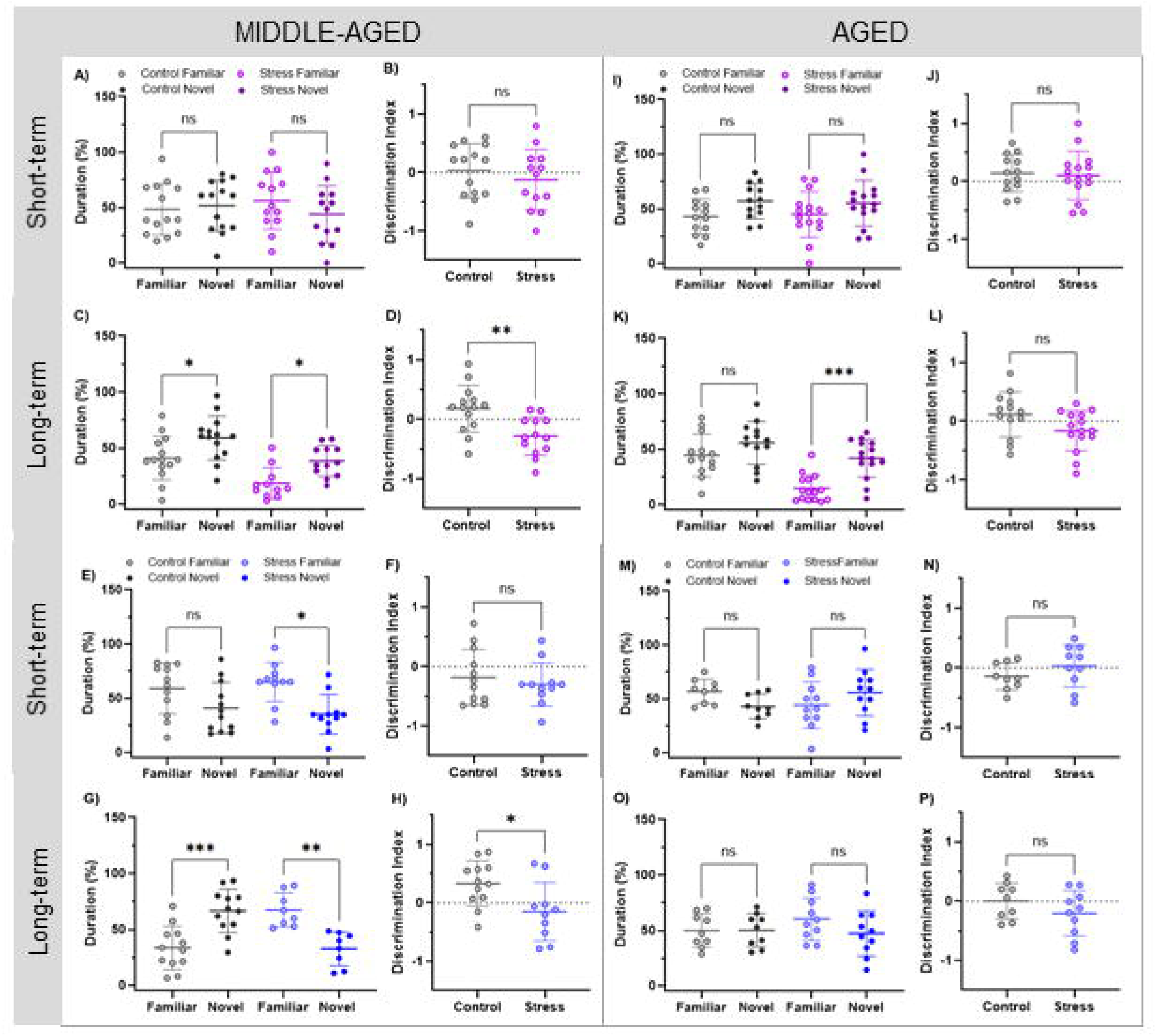
Novel object recognition performance in middle-aged and aged female (pink) and male (blue) offspring exposed to prenatal stress. Short-term memory (3 h delay) discrimination index are shown for females at 12 and 18 months (A, B, I and J) and long-term memory (24 h delay, C, D, K and L) discrimination index in the second evaluation. Male short-term memory at 12 and 18 months (E, F, M and N) and long-term memory (G, H, O and P) discrimination index. Data are presented as mean ± SD. *p < 0.05, **p < 0.01, ***p < 0.001, ns = not significant.

### 3.8. Barnes Maze Assessment

Because older control animals could not distinguish familiar from novel objects in the novel object recognition test, the Barnes maze was used to confirm and further investigate memory performance in 18-month-old offspring independently.

#### 3.8.1. Aged Females (18 Months)

- Spatial memory learning (Days 1–5): Both C and S females demonstrated reduced latency to locate the escape hole during training, indicating successful task acquisition. No significant latency differences were observed between groups during this phase (Figure 6A).
- Probe test (Day 6 – Memory check): S females spent significantly more time in the target area compared to C females (Figure 6B), indicating superior memory retention.
- Cognitive flexibility – Reversal learning (Days 7–9): Both groups showed shorter latencies during reversal learning. On Day 9, S females located the new escape hole significantly faster than C females (Figure 6C) showing enhanced cognitive flexibility.
- Probe trial (Day 10 – Cognitive flexibility assessment): No significant differences in time spent in the target quadrant were detected between groups (Figure 6D).
- Search strategy (Cognitive score): S females had lower cognitive scores, reflecting less effective search strategies, than C females during the early learning phase (Days 1–3, Figure 7). No differences were observed from days 4 to 9, indicating that S females achieved comparable search proficiency following additional training. During reversal learning, cognitive scores did not differ between groups (Figure 7).
- Approach index: C and S females did not differ significantly in the distance traveled to reach the target zone during either the memory acquisition or cognitive flexibility phases.

#### 3.8.2. Aged Males (18 Months)

- During spatial memory learning (Days 1–5), S males required significantly more time to locate the escape hole than C males on Days 3, 4, and 5 (Figure 8A), indicating impaired spatial learning. •In the Day 6 memory test, S males spent significantly less time in the target area compared to C males (Figure 8B), indicating reduced spatial memory performance.
- During the reversal learning phase (Days 7–9), both groups showed decreased completion times over the course of testing. No significant time differences were observed between groups during this phase (Figure 8C).
- On Day 10, during the mental flexibility assessment, groups did not differ significantly in time spent in the target area (Figure 8D).
- Regarding search strategy (cognitive score), groups did not differ significantly during the early learning phase (Days 1–3). However, S males exhibited significantly lower scores, reflecting less effective search strategies than C males on Days 4 and 5 (Figure 9), indicating diminished use of optimal search strategies in later learning stages. During the reversal learning phase, we detected no significant differences in scores between groups (Figure 9).
- For the approach index, S males traveled significantly greater distances to reach the target area than C males on Days 2, 3, and 5 of the memory learning phase, as well as on the final day of the mental flexibility test, indicating persistent deficits in efficient spatial navigation.

### 3.9. Assessment of Social Interaction

The three-chamber social interaction test measured the animals’ social behavior and their liking for new social partners (Figure 10). In 12-month-old females, there were no major differences in total social time or time spent in each chamber between the control (C) and experimental (S) groups during the sociability phase. However, the sociability index with the new partner (Stranger 2) was significantly lower in S females than in C females (Figure 10), indicating less interest in new social partners. At 18 months, no major differences were seen between C and S females in any measure during either the sociability or social novelty phases (Figure 10). In 12-month-old males, total social time was similar between groups during the sociability phase. The sociability index with the new partner was significantly lower in S males than in C males (Figure 10) also showing less interest in new social partners. At 18 months, no major differences were found between C and S males in any measure during either the sociability or social novelty phases (Figure 10).

**Figure 6.**
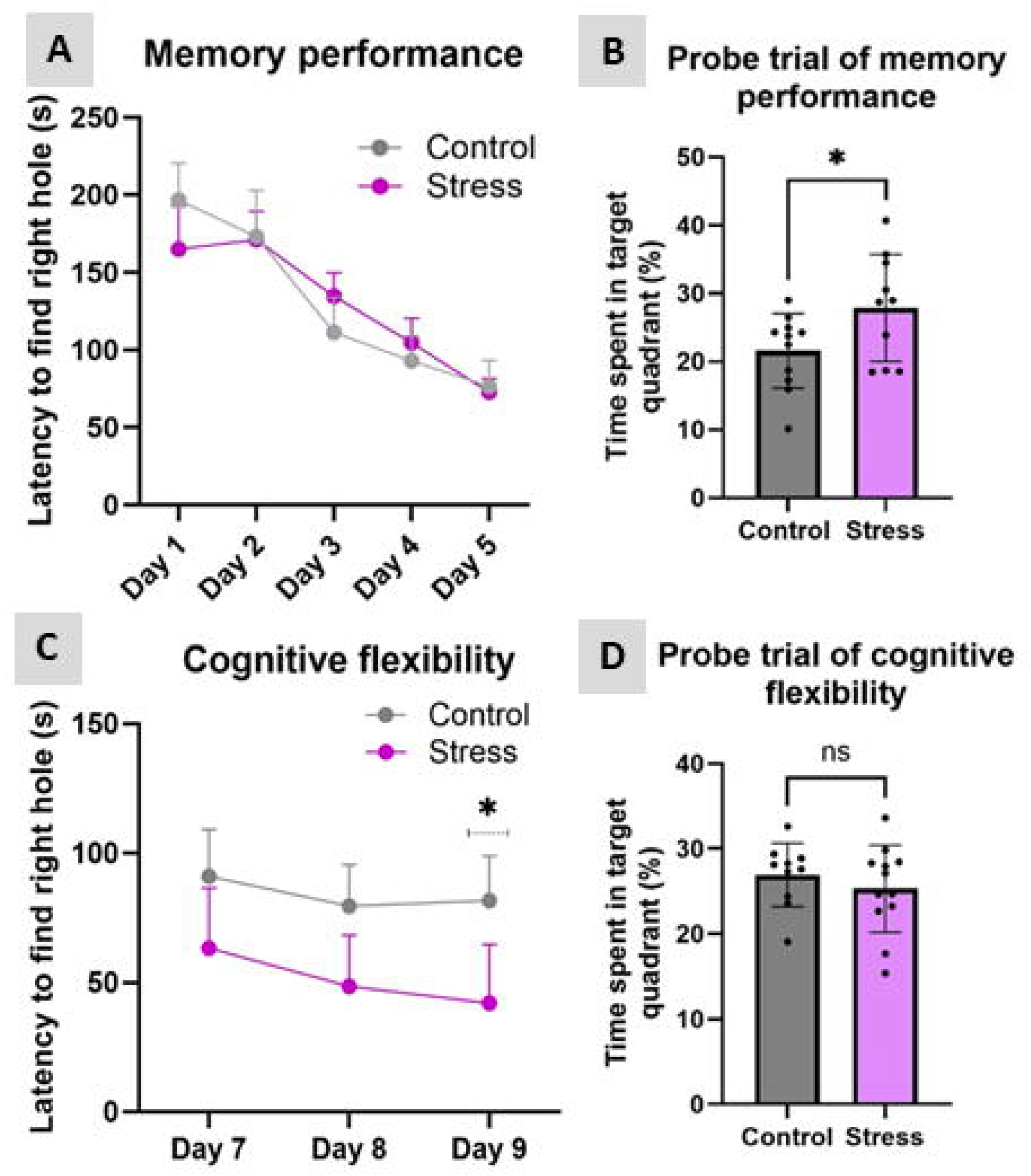
Barnes maze performance in aged female offspring exposed to prenatal stress. (A) Spatial memory acquisition phase showing the latency to locate the target hole across training days (Days 1–5). Both Control and Stress groups exhibited progressive reductions in latency over time, indicating preserved spatial learning and memory acquisition. (B) Probe trial of memory performance expressed as the percentage of time spent in the target quadrant. Females exposed to prenatal stress spent significantly more time in the target quadrant compared to Controls, suggesting enhanced spatial memory retention. (C) Cognitive flexibility assessment during reversal learning (Days 7–9). Stress animals exhibited reduced latency to locate the correct hole, with a significant difference observed on Day 9, indicating improved cognitive flexibility. (D) Probe trial of cognitive flexibility expressed as the percentage of time spent in the target quadrant. No significant differences were observed between groups. Data are presented as mean ± SD. *p < 0.05; ns = not significant.

**Figure 7.**
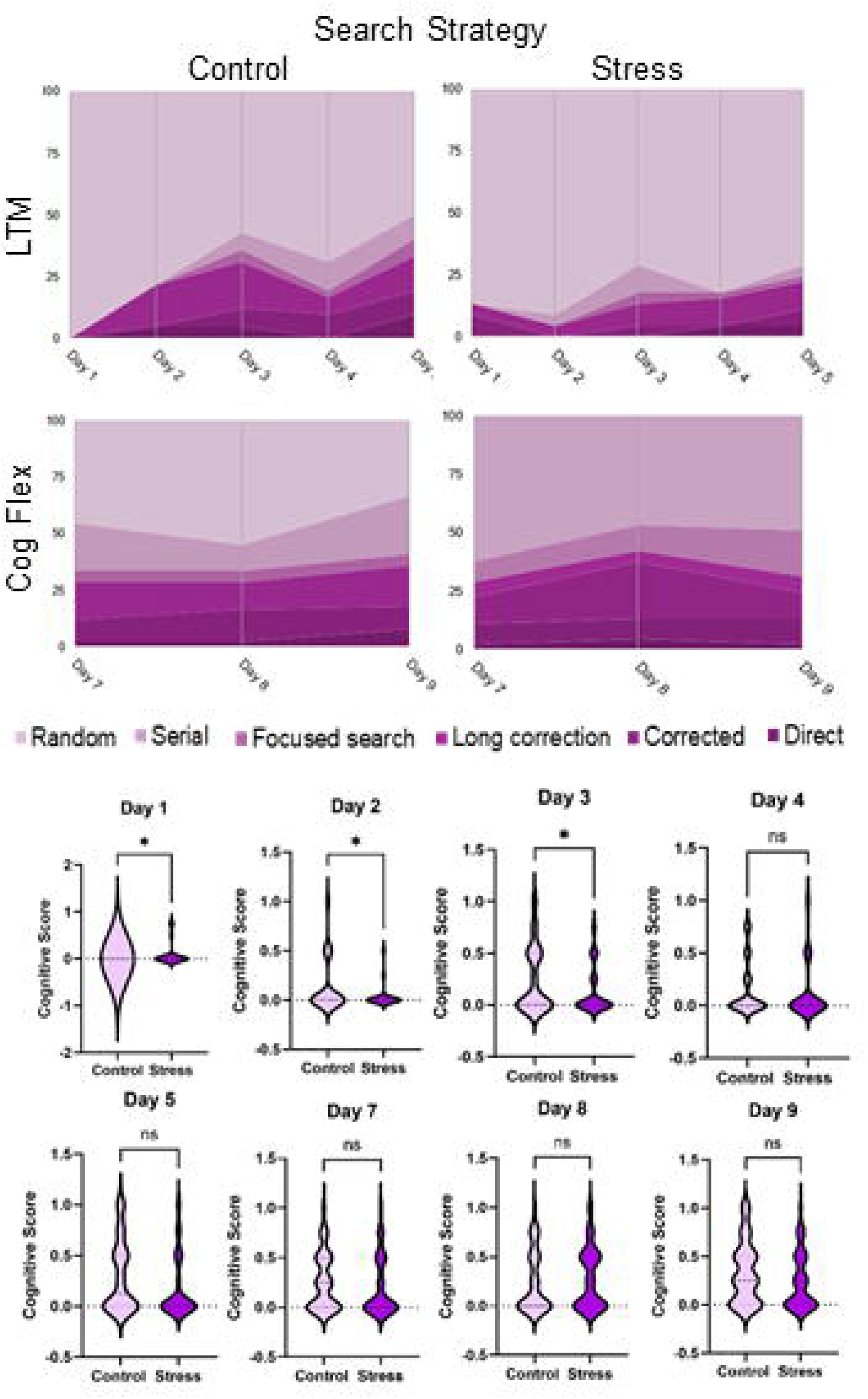
Search strategy performance in the Barnes maze of aged female offspring exposed to prenatal stress. Cognitive score during the spatial memory acquisition phase (Days 1–5). Females exposed to prenatal stress showed significantly lower cognitive scores on Days 1–3 compared to Controls, indicating impaired use of efficient search strategies during the early learning phase. No significant differences were observed on Days 4 and 5, suggesting adaptation and learning consolidation over time. Cognitive score during the cognitive flexibility/reversal learning phase (Days 7–9). No significant differences were observed between Control and Stress groups, indicating preserved search strategy performance during reversal learning. Data are presented as violin plots. *p < 0.05; ns = not significant.

**Figure 8.**
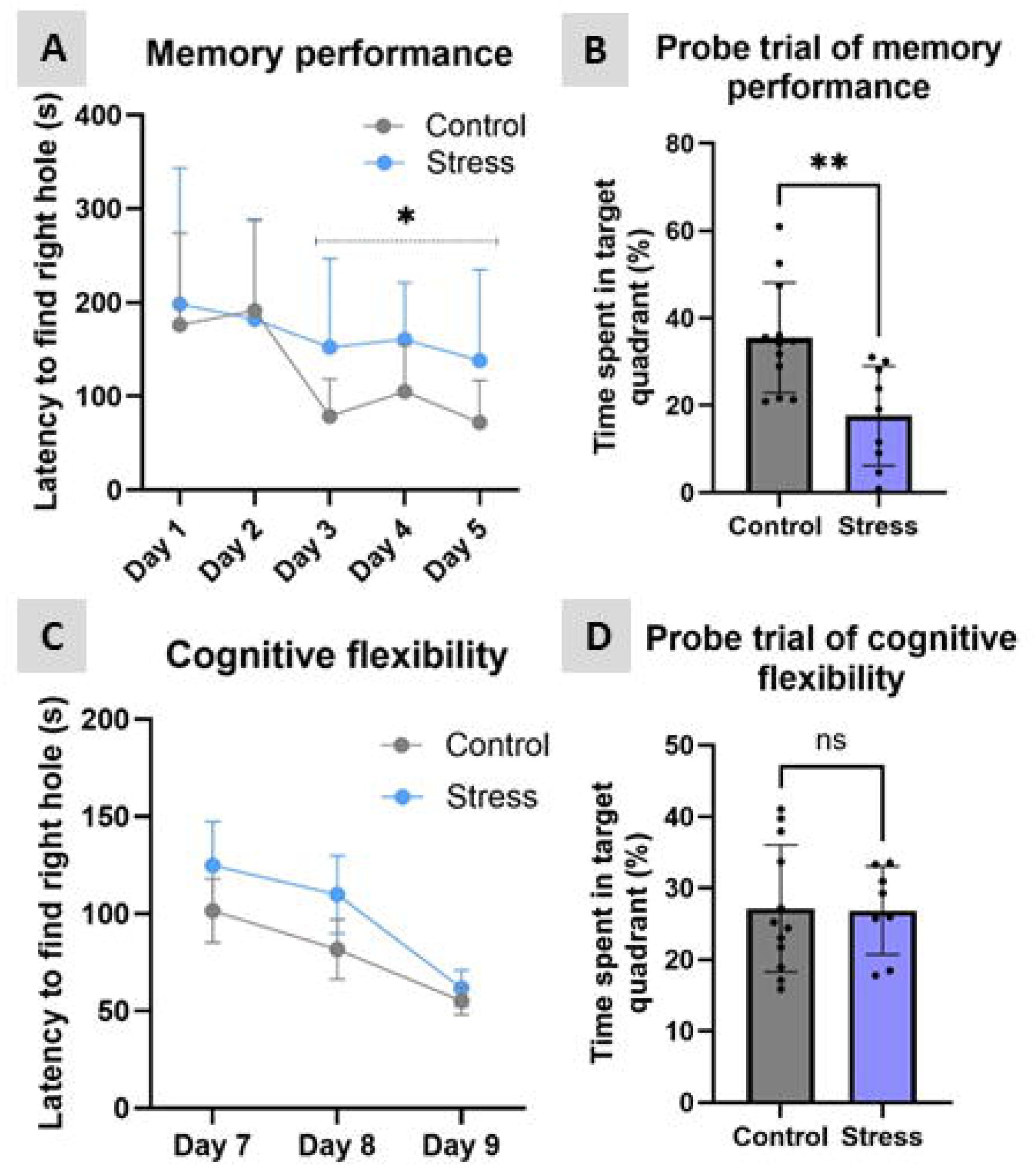
Barnes maze performance in aged male offspring exposed to prenatal stress. (A) Spatial memory acquisition phase showing the latency to locate the target hole across training days (Days 1–5). Males exposed to prenatal stress exhibited increased latency during the acquisition phase compared to Controls, indicating impaired spatial learning. (B) Probe trial of memory performance expressed as the percentage of time spent in the target quadrant. Stress animals spent significantly less time in the target quadrant than Controls, suggesting impaired spatial memory retention. (C) Cognitive flexibility assessment during reversal learning (Days 7–9). Both groups showed reduced latency across training days, indicating preserved reversal learning and cognitive flexibility. (D) Probe trial of cognitive flexibility expressed as the percentage of time spent in the target quadrant. No significant differences were observed between groups. Data are presented as mean ± SD. *p < 0.05, **p < 0.01; ns = not significant.

**Figure 9.**
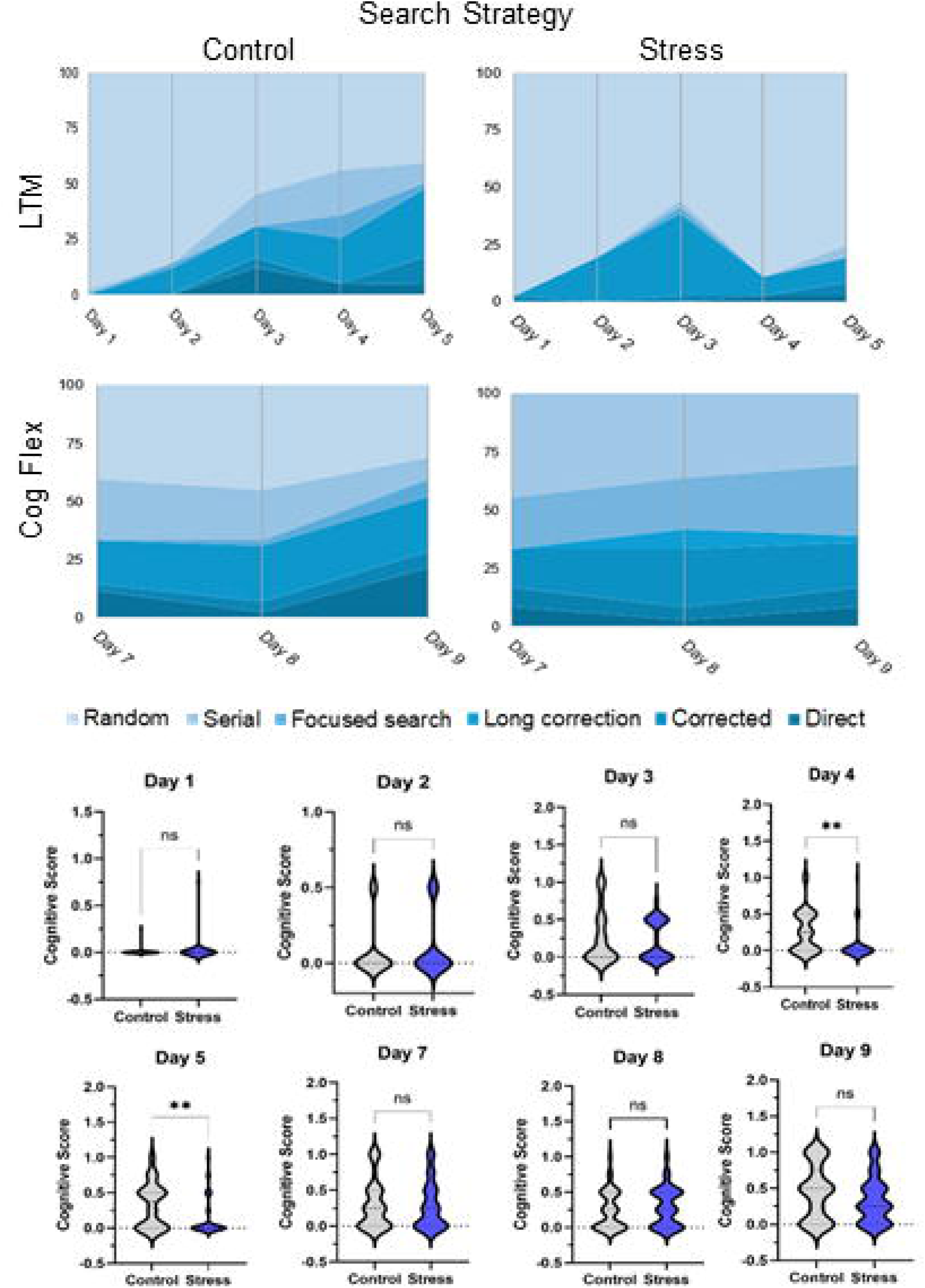
Search strategy performance in the Barnes maze of aged male offspring exposed to prenatal stress. Cognitive score during the spatial memory acquisition phase (Days 1–5). No significant differences were observed between groups during the early acquisition phase (Days 1–3). However, males exposed to prenatal stress exhibited significantly lower cognitive scores on Days 4 and 5 compared to Controls, indicating impaired use of efficient search strategies during later stages of spatial learning. Cognitive score during the cognitive flexibility/reversal learning phase (Days 7–9). No significant differences were observed between Control and Stress groups, suggesting preserved search strategy performance during reversal learning. Data are presented as violin plots. **p < 0.01; ns = not significant.

**Figure 10.**
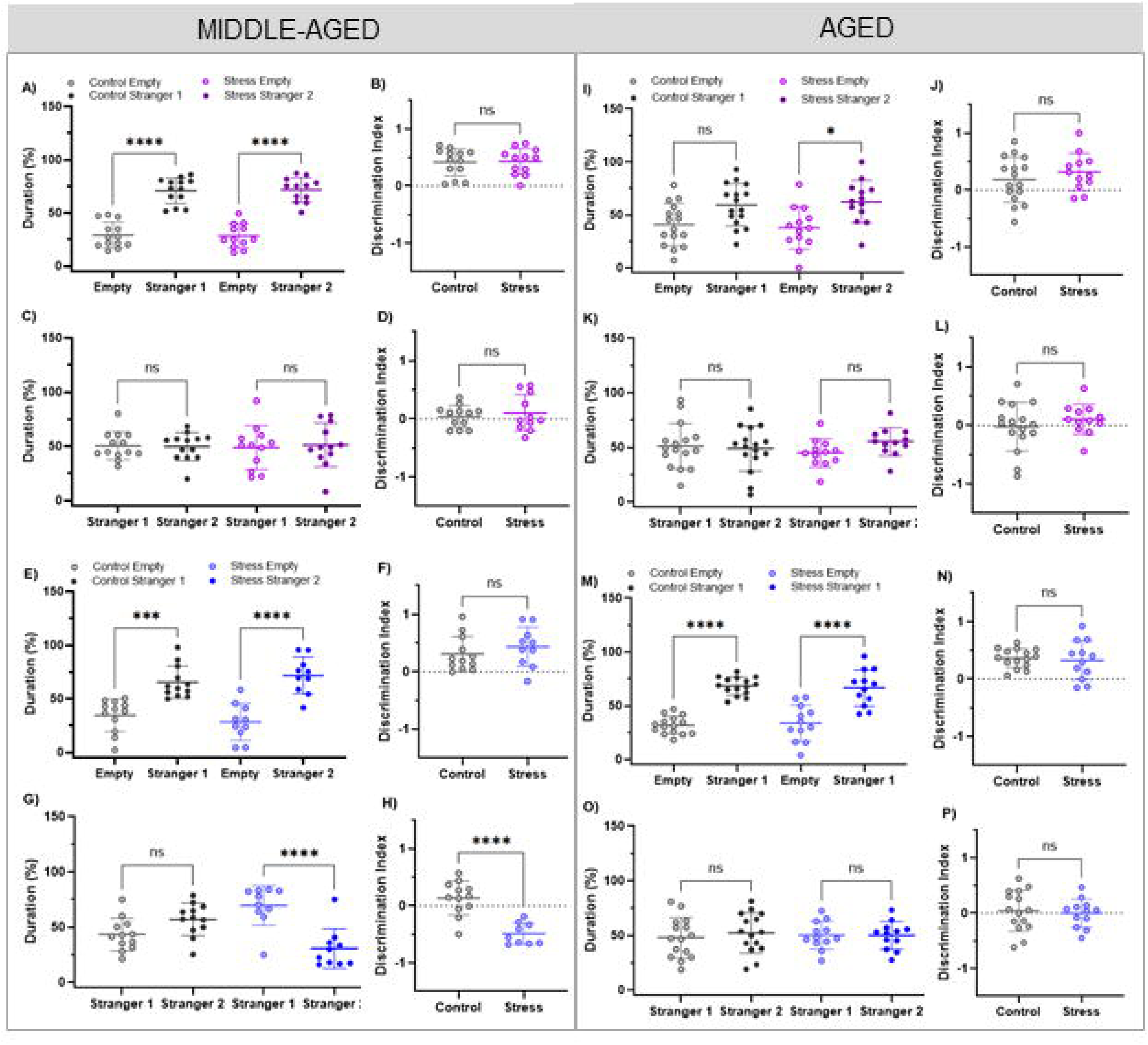
Social interaction test performance. Middle-aged female offspring exploration time (%) and discrimination index in the first evaluation (A, B). Second evaluation of exploration time of the familiar and novel stranger animals and discrimination index (C, D). Middle-aged male offspring exploration time and) discrimination index in the first evaluation (E,F). Second evaluation of middle-aged male offspring exploration time of the familiar and novel stranger animals and discrimination index (G, H). Aged females exploration and discrimination index (I, J). Exploration time and discrimination index in the second evaluation (K, L). Aged male offspring exploration time and discrimination index between groups (M, N). Exploration time and discrimination index in the second evaluation (O, P). Data are presented as mean ± SD. *p < 0.05, ***p < 0.001, ****p < 0.0001; ns = not significant.

## 4. DISCUSSION

Exposure to unpredictable mild stress (UCMS) before birth leads to persistent and distinct behavioral and cognitive trajectories in both male and female subjects across the lifespan. Early-life stress also accelerates the decline of specific brain functions, with sex-specific effects emerging from middle age (12 months) to old age (18 months). Utilizing a comprehensive battery of behavioral assessments—including measures of reward, locomotion, anxiety, memory, learning, cognitive flexibility, and social behavior—along with detailed analysis of maternal care, this study advances the understanding of fetal programming into later life stages. The behavioral data indicate that prenatal stress and unpredictable postnatal maternal care are associated with changes that shift the aging brain toward an energy-conserving state. In male rodents, this state accelerates decline in specific brain regions, while in females, it redistributes brain resources, resulting in later-life vulnerabilities in different areas. These findings clarify the influence of early life adversity on health by demonstrating that such challenges can modify both the risk of mental illness and the timing of functional decline in specific neural networks. The absence of significant changes in maternal body weight during pregnancy indicates that UCMS did not induce sustained calorie restriction during gestation. Instead, alterations in placental structure and hormone function are more plausible explanations for the observed effects. This interpretation is consistent with previous studies showing that prenatal stress can modify placental regions involved in stress hormone regulation, particularly the enzyme 11β-HSD2, and reduce placental nutrient transporter activity, thereby decreasing fetal nutrient supply without maternal weight loss (Jensen Peña et al., 2012; Briffa et al., 2017). The observed increase in stillbirths and reduced litter sizes in the stressed group suggests that UCMS adversely affects placental blood flow, increases oxidative stress, and disrupts the intrauterine environment (Bale, 2016; Zoubovsky et al., 2022). The elevated stillbirth rate may introduce survival bias and could represent a form of natural selection, potentially underestimating the full effects of prenatal stress. Offspring exposed to UCMS, regardless of sex, exhibited significantly reduced birth weights, followed by accelerated postnatal weight gain at PND7, PND14, and PND21, indicative of catch-up growth. The nutritional stress imposed by UCMS, particularly the 24-hour periods of food and water deprivation, may confound neurobehavioral outcomes. The atypical aging patterns observed in this study may result from the combined effects of excessive prenatal stress-hormone exposure and possible alterations in hippocampal insulin signaling. These findings contribute to understanding the etiology of various postnatal brain disorders. Metabolic changes can significantly affect brain health over time and are associated with increased neural damage in critical brain circuits (Wang et al., 2025). Sex differences in metabolic risk are pronounced. Male placentas exhibit a reduced capacity to upregulate 11β-HSD2 in response to maternal stress, resulting in increased glucocorticoid exposure for the male fetal brain (Bale, 2016; Bale & Epperson, 2015). This fetal exposure, combined with subsequent rapid growth, likely increases the risk of insulin resistance in offspring brains and may lay the groundwork for progressive decline in fronto-striatal brain regions, particularly in aging male offspring. In contrast, females may benefit from enhanced placental and estrogenic neural protection, potentially delaying the onset of such susceptibility. The growth trajectory identified in this study serves as an early, quantifiable marker of underlying metabolic programming and differential neural circuit vulnerabilities. Analysis of maternal behavior under unpredictable chronic mild stress (UCMS) revealed selective alterations rather than a complete loss of caregiving. Stressed mothers demonstrated reductions in licking, grooming, and nest building, while self-grooming, time spent away from the nest, and passive nursing increased. Essential maternal behaviors, including feeding, drinking, carrying pups, and active nursing, remained stable. These results suggest that stress primarily modifies the timing and type of care provided, without impairing the fundamental capacity for offspring care. This pattern of selective change aligns with previous research by Zoubovsky et al. (2020, 2022), which showed that social stress during pregnancy induces specific alterations in maternal care, leading to more fragmented and unpredictable caregiving. The present findings support the concept of maternal signal entropy, which posits that unpredictability and fragmentation contribute to adverse neurodevelopmental outcomes in childhood and adolescence, independent of the overall quantity of maternal care (Glynn et al., 2021; Baram et al., 2012; Molet et al., 2016). Howland et al. (2021) found that greater prenatal maternal mood entropy was associated with lower cognitive scores at 2 years and lower expressive language scores at 6–9 years, even after adjustment for maternal mood levels. The relatively minor disruption of nest building observed in stressed mother mice suggests a targeted effort to maintain or enhance nest organization, thereby preserving nest functionality despite broader caregiving challenges. The coexistence of both disrupted and preserved behaviors within the same individual underscores the complexity of stress effects on maternal care and suggests that chronic stress may influence distinct neural pathways governing specific maternal behaviors.

The pup retrieval test differentiated between initial maternal responsiveness and sustained caregiving. Stressed mother mice retrieved the first pup more rapidly than controls, yet the total time required to retrieve all pups remained similar across groups. These results suggest that stressed mothers display heightened initial alertness and responsiveness, followed by diminished attentiveness during subsequent retrievals. In developmental neuroscience, unpredictability in maternal signals is regarded not only as inadequate care but also as a programming signal that can disrupt the maturation of neural circuits in newborns (Baram et al., 2012; Tottenham, 2020). The altered maternal behavior observed here may reduce the predictability of neonatal cues. Maternal and environmental signals contribute to early circadian development (Wong et al., 2022), providing a possible link between disrupted caregiving and later behavioral outcomes. In this model, circadian disruption is a proposed developmental mechanism rather than a measured mediator of brain aging.

The accelerated aging observed in male mice may demonstrate how neural timing mechanisms are affected by the combined influence of prenatal stress hormone exposure and reduced placental 11β-HSD2 activity (Bale, 2016). Adverse prenatal conditions and postnatal unpredictability likely represent interconnected processes that shape the trajectory of brain aging.

Behavioral patterns in stressed females at two distinct time points demonstrate initial disturbances, followed by the preservation of specific cognitive functions. This observation aligns with the hypothesis that the brain reallocates resources under energy constraints. At 12 months, stressed females displayed reduced locomotion, decreased interest in novel social contexts, and lower anxiety-like behaviors. These findings likely result from alterations in dopaminergic and oxytocinergic signaling pathways, which may contribute to reduced motivation and modified responses to social reward (Bäckman et al., 2010; Russo & Nestler, 2013; Hurlemann & Grinevich, 2018).

The observed reduction in anxiety-like behavior among middle-aged, stressed females may indicate alterations in the hypothalamic-pituitary-adrenal (HPA) axis following repeated prenatal stress (Heck & Handa, 2019; Babb et al., 2014). Supporting evidence suggests that chronic stress exposure decreases HPA axis sensitivity, as indicated by reduced corticotropin-releasing hormone (CRH) receptor expression and diminished corticosterone release in response to novel stressors (McEwen & Akil, 2020; Babb et al., 2014). These neurobiological adaptations are thought to be mediated by epigenetic mechanisms, including increased methylation of CRH-related genes and elevated levels of brain-derived neurotrophic factor (BDNF) and GABAergic components (Roth et al., 2009; Weaver et al., 2004; Szyf, 2013). However, the increased time stressed females spent in the open arms of the elevated plus maze (EPM) may instead indicate impaired risk assessment or diminished behavioral regulation, particularly given their reduced locomotion in the open field test (Walf & Frye, 2007).

In stressed female rodents, unlike males, the transition from midlife to old age is marked by the onset of late-stage anhedonia, while hippocampus-dependent memory remains normal. Thus, anhedonia is not seen at 12 months, suggesting a decline in dopamine-dependent brain circuits and providing evidence that estrogen may regulate dopamine function to prevent anhedonic behavior. However, as estrogen levels drop during the shift to reproductive aging, this protection weakens (Becker, 1999; Yagi & Galea, 2019; Frick & Kim, 2018). The enhanced memory performance observed in stressed, aged females represents a notable finding of this study. However, this outcome should not be interpreted as cognitive enhancement or simple resilience. Instead, it aligns with a pattern of neural resource reallocation under allostatic constraints (McEwen & Gianaros, 2011). In brains subjected to early-life adversity, homeostatic mechanisms may prioritize preserving circuits for immediate survival, such as those involved in spatial navigation and hippocampal memory, over others, including the dopaminergic pathway. At 18 months, preserved spatial memory coexisted with reduced sucrose preference, an operational measure of anhedonia-like behavior. This dissociation suggests different trajectories of memory and reward-related behavior. Estrogen-dependent hippocampal plasticity (Frick & Kim, 2018) and reward-circuit mechanisms of anhedonia (Russo & Nestler, 2013) provide complementary frameworks for interpreting these trajectories. We use “compensatory resilience” to describe selective preservation of memory despite impairment in another behavioral domain; resource reallocation and its dopaminergic basis remain mechanistic interpretations rather than demonstrated causes.

Findings from the novel object recognition (NOR) and Barnes maze tests demonstrate that stressed female subjects maintain hippocampus-dependent memory function and exhibit enhanced cognitive flexibility during reversal learning at 18 months. The pattern of initial delay followed by subsequent improvement suggests that compensatory mechanisms primarily support memory storage rather than acquisition. Furthermore, early learning deficits likely reflect prefrontal cortex dysfunction rather than hippocampal impairment (Nyberg et al., 2012; Cabeza et al., 2018). Experimental and review literature links BDNF–TrkB signaling, CREB-dependent plasticity, dendritic remodeling, and estrogen-dependent hippocampal function to cognitive performance. These mechanisms offer possible explanations for the behavioral preservation observed here (Zhou et al., 2023; Faraji & Metz, 2025; Olave et al., 2022; Serita et al., 2017; Frick & Kim, 2018). Baumgartner et al. (2022) showed that short-term estradiol exposure during midlife in female rodents induces persistent alterations in hippocampal estrogen synthesis. Interactions between local estrogen synthesis and IGF-1 signaling can favor ERK/MAPK-dependent phosphorylation of estrogen receptor alpha (ERα) at serine 118 and sustained receptor activity (Baumgartner et al., 2022; Flannery et al., 2025). Evidence that neuroestrogens regulate hippocampal parvalbumin interneurons and their perineuronal nets (Hernández-Vivanco et al., 2025) further supports this candidate explanation for sex differences in memory preservation.

In contrast, the mesolimbic dopaminergic system lacks the local estrogen-IGF-1-Erα cycle, as hippocampal aromatase expression relies on circulating estrogens (Ma et al., 2020; Chen et al., 2021). Although overall memory performance remained intact, aged and stressed female offspring demonstrated less efficient search strategies during the initial phase of Barnes maze acquisition. This finding indicates that prenatal stress and aging may specifically impact prefrontal cortical circuits. Prenatal stress may induce an excitatory-inhibitory imbalance in the prefrontal cortex, disrupt glutamatergic gene expression, reduce inhibitory transmission, limit synapse formation, and decrease prefrontal cortex adaptability (Holmes & Wellman, 2009). These stress-induced alterations may interact with intrinsic aging processes, resulting in prefrontal hypofunction (Zohar & Weinstock, 2011). The observed behavioral profile, characterized by preserved hippocampal memory but impaired prefrontal strategy, suggests a dissociation between hippocampal preservation and prefrontal vulnerability. This pattern is consistent with the differential sensitivity of these neural circuits to glucocorticoid exposure, prenatal programming, and biological aging (McEwen & Morrison, 2013; Spellman et al., 2015). Conversely, maternal prenatal stress in male offspring appears to initiate a gradual decline, beginning with early dysfunction in the mesolimbic region and progressing to widespread impairments in the hippocampal-prefrontal brain areas. In midlife, stressed males exhibit reduced sucrose preference, a recognized indicator of anhedonia and dopamine system impairment. This phenotype is associated with alterations in dopamine D2 receptor expression in the nucleus accumbens and prefrontal cortex, which may accelerate aging in reward-related neural circuits (Russo & Nestler, 2013; Berridge & Kringelbach, 2015; Mohebi et al., 2019; Weinstock, 2017).

Here, as Mohebi et al. (2019) observed, reduced jumping behavior among stressed middle-aged males in the open field, without other movement alterations, may indicate a specific decrease in upward exploratory drive, potentially linked to diminished dopamine-mediated motivation. Behavioral assessments across multiple tests provided clear evidence of sex-specific temporal divergence in anxiety-like behavior during aging. Middle-aged males did not exhibit a significant decrease in anxiety-like behavior, whereas females demonstrated a reduction. At 18 months, stressed males entered old age, characterized by a transition from low movement to increased activity. This shift included greater open field activity, such as increased distance traveled, higher speed, more frequent walking, rearing, jumping, and decreased resting. These behavioral changes were associated with reduced anxiety-like behavior but were also accompanied by persistent spatial memory deficits. The coexistence of increased activity and persistent memory deficits is compatible with altered regulation of motor and cognitive behavior during aging. Age-related changes in dopaminergic function provide a plausible context for this interpretation (Collier et al., 2011; Bäckman et al., 2010). The behavioral pattern suggests fronto-striatal dysfunction, without establishing dopaminergic neuron loss or a compensatory response to that loss.

In male offspring, the lack of estrogen-mediated maintenance of perineuronal nets around parvalbumin interneurons, as observed in female mice (Hernández-Vivanco et al., 2025), may lead to persistent inflammatory changes in the extracellular matrix. These changes could progressively impair fast-spiking inhibitory interneurons, which are essential for neural circuit regulation. Impaired gamma-aminobutyric acid (GABA) clearance is hypothesized to contribute to the excessive motor activity observed in older male offspring. Although these molecular mechanisms are inferred from previous studies, direct validation through neurochemical analyses is still needed.

Persistent spatial memory deficits observed in older male offspring result from the combined detrimental effects of hormonal imbalance on the brain. In contrast, females exhibit greater neural hormonal adaptability, as previously discussed, whereas males lack this degree of humoral protection. Consequently, the male hippocampus is especially susceptible to progressive dysfunction caused by dopamine depletion associated with rapid growth, ultimately contributing to cognitive decline (Wang et al., 2025; Grillo et al., 2019). Within this context, increased exploratory behavior in stressed male offspring does not represent adaptive exploration. Rather, it indicates an uncontrolled drive to move and persistent, ineffective random searching associated with dysfunction in fronto-striatal brain circuits. The observed increase in rearing and jumping behaviors is unlikely to reflect heightened curiosity and may instead result from repetitive motor actions driven by dopamine dysregulation and excessive neural activity. This interpretation positions the behavior of older stressed male offspring within the broader context of accelerated aging in fronto-striatal brain regions. A diminished regulatory influence of the prefrontal cortex on other brain regions, leading to decreased fear-based avoidance and impaired movement control, produces behavior that superficially resembles reduced anxiety in the elevated plus maze (EPM) but is mechanistically distinct. In contrast, genuine anxiety reduction would preserve cognitive capacity and effective exploratory strategies. The observed combination of decreased avoidance and poor strategic movement instead reflects a broader loss of executive control associated with reduced prefrontal cortical function (McEwen & Morrison, 2013; Dalley & Robbins, 2017; Bäckman et al., 2010).

We repeatedly observed significant, consistent temporal differences in search strategy performance between male and female progeny. Offspring of gestationally stressed females employed less effective strategies during the initial learning phase (Days 1 to 3); however, their performance improved to match control levels by Days 4 to 5, indicating delayed adaptation. In contrast, stressed male offspring demonstrated effective strategies during early learning, but their performance declined during Days 4 to 5, suggesting impaired retention of previously acquired knowledge. This pattern aligns with the NOR results, which indicate that prenatal stress differentially influences learning and memory processes by sex. Female progeny exhibit early learning deficits but retain memory later, whereas males display effective initial learning but impaired memory retention. The observed memory retention deficits in male progeny may result from structural and functional changes in the hippocampus.

This study corroborates previous findings that both sexes retain adaptive abilities during reversal learning, although male offspring show impaired spatial memory. These findings suggest that prenatal stress selectively impairs hippocampus-dependent memory while largely preserving prefrontal cortex-mediated cognitive flexibility. This distinction is consistent with evidence that the hippocampus and prefrontal cortex respond differently to hormonal influences during development (McEwen & Morrison, 2013), supporting the view that gestational stress affects cognition within specific neural domains rather than uniformly across brain functions. Reduced testosterone levels or androgen-related dysfunctions may further contribute to motivational decline in males. Several studies have demonstrated that gestational stress disrupts the hypothalamic-pituitary-gonadal axis, resulting in persistent impairments in gonadal hormone production, including testosterone (Toufexis et al., 2014). The combined effects of declining gonadal hormones and established dopaminergic vulnerability may exacerbate deficits in reward and motivational circuits in aged males. A specific reduction in interest in novel social partners was observed at 12 months in both male and female mice, while overall social interaction remained unchanged at 18 months. The 12-month results indicate a selective deficit in motivation for novel social experiences, rather than a general impairment in social behavior. Social reward involves distributed circuits (Matyjek et al., 2020), including a medial amygdala–preoptic pathway that promotes accumbal dopamine release (Hu et al., 2021) and coordinated accumbal oxytocin–serotonin signaling (Dölen et al., 2013). We interpret reduced social novelty preference in relation to these reward mechanisms. The apparent improvement at 18 months, however, may reflect disorganized social behaviors in males, potentially increasing social interaction and masking underlying difficulties in recognizing novel social contexts. In females, alterations in energy metabolism and delayed onset of anhedonia may modify typical social motivation. These age-dependent findings could be confounded by variability in sample sizes over time, rather than reflecting complete recovery. Although all testing was conducted during the estrous phase, evidence suggests that even within a specific stage, residual hormonal fluctuations and uncontrolled variability in dopamine and oxytocin may significantly influence social interaction behavior in female mice (Chari et al., 2020).

Current findings suggest that sex differences in brain aging trajectories begin during fetal development (Bale, 2016; Bale & Epperson, 2015), when adverse prenatal effects may differ by sex and time and influence neural connectivity (Islam et al., 2022; Georgousopoulou et al., 2026). These sex- and time-based differences suggest that prenatal stress differentially programs the timing of functional decline in distinct brain circuits. In males, prenatal stress is posited to induce early vulnerability in the mesolimbic dopaminergic pathway, which manifests as anhedonia during midlife. As aging progresses, this initial dopaminergic vulnerability is hypothesized to extend to fronto-striatal and hippocampal-prefrontal circuits. Sex-dependent estrogenic regulation of hippocampal parvalbumin interneurons and their perineuronal nets (Hernández-Vivanco et al., 2025) suggests that extracellular matrix plasticity could contribute to divergent cognitive trajectories. Extension of this mechanism to the male medial prefrontal cortex and dorsal striatum is a component of the proposed model. At the same time, catch-up growth may induce central insulin resistance, impairing the IRS-1/Akt signaling pathway and compromising synaptic plasticity (Grillo et al., 2019; Wang et al., 2025). The convergence of perineuronal metabolic dysfunction and chronic neuroinflammation is hypothesized to precipitate a failure of top-down inhibitory control. This failure may result in persistent spatial memory deficits, impaired search strategies, motor perseveration, and a global disinhibitory syndrome. This trajectory suggests progressive and cumulative deterioration across multiple neural domains, consistent with accelerated fronto-striatal decline. However, certain hippocampal circuits exhibit resilience in old age. A candidate mechanism involves interactions between hippocampal neuroestrogens, IGF-1, and ERα signaling that can sustain plasticity beyond an initial estradiol exposure (Baumgartner et al., 2022; Flannery et al., 2025). The coexistence of preserved memory and reduced sucrose preference motivates an allostatic compensation hypothesis. It does not establish that energy is transferred from reward circuits to the hippocampus. Age-related changes in estrogen signaling and neuroinflammation may modify the balance between hippocampal plasticity and reward-related behavior (Lephart et al., 2026). This provides a possible explanation for the late emergence of anhedonia-like behavior in females. Unpredictable maternal signals have been linked to later neurobehavioral outcomes (Baram et al., 2012; Molet et al., 2016), while maternal cues contribute to early circadian development (Wong et al., 2022). Together, these findings support a developmental programming hypothesis for the age-dependent behavioral changes observed here. In males, whose placentas exhibit reduced 11β-HSD2 protective capacity (Bale, 2016), increased glucocorticoid exposure combined with postnatal signal unpredictability may accelerate functional aging and result in a progressive deterioration phenotype.

In contrast, placental protection during gestation and estrogen-dependent hippocampal plasticity later in life may contribute through distinct developmental stages to the female behavioral trajectory. Preservation of memory alongside anhedonia-like behavior suggests that these effects vary across functional domains. The proposed model, integrating the UCMS protocol with accelerated postnatal catch-up growth, likely establishes a synergistic programming axis in which central insulin resistance and glucocorticoid neurotoxicity converge on hippocampal and prefrontal circuits (Grillo et al., 2019). This neuroendocrine-metabolic continuum may explain the progressive and cumulative cognitive decline observed in males, as metabolic dysfunction exacerbates the initial glucocorticoid-induced vulnerability across the lifespan.

Negative findings, such as the lack of anxiety changes in middle-aged males or the absence of social interaction deficits at 18 months, may indicate either a true absence of effect or insufficient statistical power to detect moderate differences (Button et al., 2013; Lazic, 2010). Therefore, this model should be considered a preliminary framework that requires further validation through rigorously designed studies with adequate sample sizes, control for litter effects, inclusion of additional time points to clarify the timing of circuit vulnerabilities, and direct molecular and protein-level analyses to confirm the proposed neural circuit mechanisms. These findings emphasize the necessity of including sex as a biological variable in studies of fetal development and brain aging, and highlight the need for sex-specific strategies in the prevention and treatment of stress-related brain disorders throughout the lifespan. Identifying critical transitional periods, such as midlife in males and the perimenopausal period in females, may inform the optimal timing of preventive interventions to mitigate the long-term effects of prenatal stress.

These findings indicate that UCMS experienced by mothers induces distinct behavioral and cognitive trajectories in males and females across the lifespan, with unique alterations emerging from middle age to old age. In males, accelerated degeneration of the fronto-striatal brain region is observed, likely resulting from stress-related hormonal damage and increased rigidity in the medial prefrontal cortex due to reduced local estrogen activity. These neurobiological changes contribute to persistent deficits in spatial memory, repetitive behaviors, and impaired risk assessment. The male trajectory shows a gradual decline across multiple brain regions, suggesting progressive depletion of neural resources that warrants further investigation.

In contrast, females exhibit a pattern termed allostatic reallocation. During midlife, modifications in neural circuits associated with anxiety and motor function compensate for vulnerabilities in the mesolimbic pathway. In old age, a self-sustaining molecular cycle, potentially involving hormonal fluctuations, appears to preserve hippocampus-dependent memory through transient estrogen-mediated mechanisms. This adaptive process represents a trade-off, reallocating resources from mesolimbic dopamine activity to sustain navigational abilities. However, the decline in estrogen during reproductive aging may ultimately disrupt this protective mechanism, triggering neuroinflammation and resulting in late-onset anhedonia. Consequently, the mesolimbic trade-off constrains both the duration and efficacy of this compensatory strategy. Temporal differences in behavioral outcomes suggest that maternal prenatal stress combined with unpredictable postnatal care jointly influence sex-specific trajectories of brain aging. Epigenetic mechanisms may contribute, and sex differences in placental stress hormone responses are evident. Male placentas demonstrate reduced capacity to initiate protective responses, resulting in substantially higher stress hormone exposure for the developing male brain. These findings underscore the importance of considering sex as a critical biological variable in studies of fetal development. Identifying key transitional periods, such as midlife in males and the perimenopausal phase in females, may inform targeted prevention strategies. While certain experimental limitations exist, they do not diminish the results’ significance. Although the unpredictable chronic mild stress (UCMS) paradigm is well established in prenatal stress research, we did not directly measure maternal corticosterone levels. Nevertheless, reduced litter size and birth weight provide indirect evidence of effective glucocorticoid exposure. Additionally, we infer the association of behavioral phenotypes with specific neural circuit dysfunctions, such as the estrogen-IGF-1-estrogen receptor alpha loop, perineuronal net stiffening, or impaired GABAergic gating, from prior behavioral studies. Including metabolic stressors in the UCMS protocol adds complexity to the underlying mechanisms. Unequal group sizes limited the statistical power to detect moderate effects in certain comparisons. Administering stress across the entire gestational period precludes identification of critical windows of vulnerability for specific neural circuits. Furthermore, the increased stillbirth rate observed in the stressed group may introduce survival bias. Future investigations should incorporate direct molecular and neurochemical assessments, cross-fostering designs, epigenetic analyses, and comprehensive metabolic profiling to elucidate the mechanisms underlying sex-specific patterns of brain aging. Proteomic studies are also necessary to validate the proposed hyperactive signaling pathways in females and to clarify the molecular basis of fronto-striatal dysfunction in males.

## 5. CONCLUSIONS

In summary, this study demonstrates that exposure to unpredictable chronic mild stress (UCMS) during prenatal life, combined with fragmented postnatal maternal care, establishes a synergistic neuroendocrine and metabolic programming continuum that shapes sexually divergent trajectories of brain aging in C57BL/6J mice. Our findings reveal distinct temporal and region-specific circuit vulnerabilities between sexes that manifest progressively from midlife to old age:

- In males, prenatal adversity initiates a continuous, multi-domain cognitive and behavioral decline. This trajectory begins with early mesolimbic dopaminergic dysfunction (evident as midlife anhedonia) and culminates in old age as an accelerated fronto-striatal decline. This advanced stage is characterized by marked spatial memory deficits, poor strategic navigation, motor perseveration, and a global disinhibitory syndrome, likely aggravated by prenatal glucocorticoid overexposure, postnatal catch-up growth, and localized insulin resistance.
- In females, the aging brain exhibits a pattern of allostatic reallocation. Stressed female offspring demonstrate a transient, cell-autonomous preservation of hippocampal plasticity and navigational competence that lasts into old age. This specific cognitive resilience is supported by local neuroestrogen- and growth factor-dependent pathways. However, this adaptive mechanism comes at a clear trade-off, progressively compromising dopamine-dependent reward circuitry and precipitating late-onset anhedonia as circulating estrogens decline during reproductive senescence.

Importantly, the preservation of cognitive flexibility during reversal learning in both sexes indicates domain-specific rather than global cognitive deterioration. Ultimately, these insights underscore the critical necessity of incorporating sex as a biological variable in fetal programming and biogerontology research across the lifespan. Identifying these sex-specific trajectories and their critical transitional windows—such as midlife for males and reproductive senescence for females— will be fundamental for developing targeted, timely preventive and therapeutic interventions against stress-induced neurodegenerative phenotypes.

## 6. Declaration of generative ai and ai-assisted technologies in the manuscript preparation process

During the preparation of this work the author(s) used Claude Opus 5 in order to correlate the large number of results. After using this tool/service, the author(s) reviewed and edited the content as needed and take(s) full responsibility for the content of the published article. The graphical abstract was designed with the assistance of OpenAI’s ChatGPT (GPT-5.6) for visual conceptualization and illustration. All scientific content, interpretation, and final figure design were reviewed and approved by the authors.

## 7. Funding statement

This research was partially funded by the Coordenação de Aperfeiçoamento de Pessoal de Nível Superior (CAPES), the National Council for Scientific and Technological Development (CNPq) [308043/2022-7], Brazilian National Institute of Science and Technology of DOHaD-INCT [409165/2024-7], São Paulo Research Fundation (FAPESP) [2024/06071-1].

---

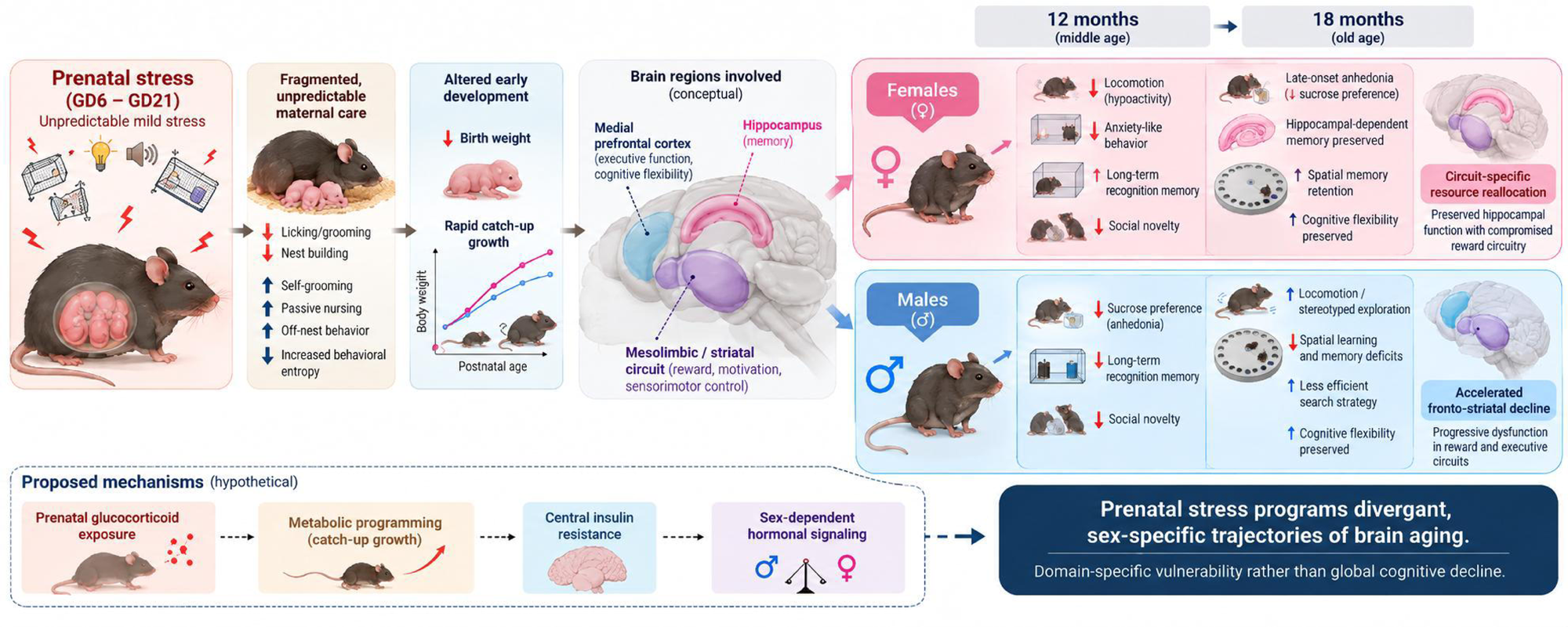

## Notes

### Competing Interest Statement

The authors have declared no competing interest.

## REFERENCES

1. Antunes, M., & Biala, G. (2012). The novel object recognition memory: neurobiology, test procedure, and its modifications. Cognitive Processing, 13(2), 93–110. 10.1007/s10339-011-0430-z

2. Babb, J. A., Masini, C. V., Day, H. E. W., & Campeau, S. (2014). Habituation of hypothalamic– pituitary–adrenocortical axis hormones to repeated homotypic stress and subsequent heterotypic stressor exposure in male and female rats. Stress, 17(3), 224–234. 10.3109/10253890.2014.905534

3. Bäckman, L., Lindenberger, U., Li, S. C., & Nyberg, L. (2010). Linking cognitive aging to alterations in dopamine neurotransmitter functioning: Recent data and future avenues. Neuroscience & Biobehavioral Reviews, 34(5), 670–677. 10.1016/j.neubiorev.2009.12.008

4. Bale, T. L. (2016). The placenta and neurodevelopment: sex differences in prenatal vulnerability. Dialogues in Clinical Neuroscience, 18(4), 459–464. 10.31887/dcns.2016.18.4/tbale

5. Bale, T. L., & Epperson, C. N. (2015). Sex differences and stress across the lifespan. Nature Neuroscience, 18(10), 1413–1420. 10.1038/nn.4112

6. Baram, T. Z., Davis, E. P., Obenaus, A., Sandman, C. A., Small, S. L., Solodkin, A., et al. (2012). Fragmentation and Unpredictability of Early-Life Experience in Mental Disorders. American Journal of Psychiatry, 169(9), 907–915. 10.1176/appi.ajp.2012.11091347

7. Barker, D. J. P. (Ed.) (1992). Fetal and infant origins of adult disease. BMJ Books.

8. Barker, D. J. P. (1994). Maternal and Fetal Origins of Coronary Heart Disease. Journal of the Royal College of Physicians of London, 28(6), 544–551. 10.1016/s0035-8819(25)00700-7

9. Barker, D. J. P. (1995). Fetal origins of coronary heart disease. BMJ, 311(6998), 171–174. 10.1136/bmj.311.6998.171

10. Baumgartner, N. E., McQuillen, S. M., Perry, S. F., Miller, S., Maroteaux, M. J., Gibbs, R. B., et al. (2022). History of Previous Midlife Estradiol Treatment Permanently Alters Interactions of Brain Insulin-like Growth Factor-1 Signaling and Hippocampal Estrogen Synthesis to Enhance Cognitive Aging in a Rat Model of Menopause. Journal of Neuroscience, 42(42), 7969–7983. 10.1523/jneurosci.0588-22.2022

11. Becker, J. B. (1999). Gender Differences in Dopaminergic Function in Striatum and Nucleus Accumbens. Pharmacology Biochemistry and Behavior, 64(4), 803–812. 10.1016/s0091-3057(99)00168-9

12. Benoit, J. D., Rakic, P., & Frick, K. M. (2015). Prenatal stress induces spatial memory deficits and epigenetic changes in the hippocampus indicative of heterochromatin formation and reduced gene expression. Behavioural Brain Research, 281, 1–8. 10.1016/j.bbr.2014.12.001

13. Berridge, K. C., & Kringelbach, M. L. (2015). Pleasure Systems in the Brain. Neuron, 86(3), 646–664. 10.1016/j.neuron.2015.02.018

14. Berrio, J. P., Hestehave, S., & Kalliokoski, O. (2024). Reliability of sucrose preference testing following short or no food and water deprivation—a Systematic Review and Meta-Analysis of rat models of chronic unpredictable stress. Translational Psychiatry, 14(1), 39. 10.1038/s41398-024-02742-0

15. Briffa, J. F., Hosseini, S. S., Tran, M., Moritz, K. M., Cuffe, J. S. M., & Wlodek, M. E. (2017). Maternal growth restriction and stress exposure in rats differentially alters expression of components of the placental glucocorticoid barrier and nutrient transporters. Placenta, 59, 30–38. 10.1016/j.placenta.2017.09.006

16. Button, K. S., Ioannidis, J. P. A., Mokrysz, C., Nosek, B. A., Flint, J., Robinson, E. S. J., et al. (2013). Power failure: why small sample size undermines the reliability of neuroscience. Nature Reviews Neuroscience, 14(5), 365–376. 10.1038/nrn3475

17. Cabeza, R., Albert, M., Belleville, S., Craik, F. I. M., Duarte, A., Grady, C. L., et al. (2018). Maintenance, reserve and compensation: the cognitive neuroscience of healthy ageing. Nature Reviews Neuroscience, 19(11), 701–710. 10.1038/s41583-018-0068-2

18. Chari, T., Griswold, S., Andrews, N. A., & Fagiolini, M. (2020). The Stage of the Estrus Cycle Is Critical for Interpretation of Female Mouse Social Interaction Behavior. Frontiers in Behavioral Neuroscience, 14, 113. 10.3389/fnbeh.2020.00113

19. Chen, H. B., Xu, C., Zhou, M. H., Qiao, H., & An, S. C. (2021). Endogenous hippocampal, not peripheral, estradiol is the key factor affecting the novel object recognition abilities of female rats. Behavioral Neuroscience, 135(5), 668–679. 10.1037/bne0000480

20. Collier, T. J., Kanaan, N. M., & Kordower, J. H. (2011). Ageing as a primary risk factor for Parkinson’s disease: evidence from studies of non-human primates. Nature Reviews Neuroscience, 12(6), 359–366. 10.1038/nrn3039

21. Dalley, J. W., & Robbins, T. W. (2017). Fractionating impulsivity: neuropsychiatric implications. Nature Reviews Neuroscience, 18(3), 158–171. 10.1038/nrn.2017.8

22. de Sousa Júnior, G. M., Tavares, V. D. O., de Meiroz Grilo, M. L. P., Coelho, M. L. G., de Lima-Araújo, G. L., Schuch, F. B., et al. (2021). Mental Health in COVID-19 Pandemic: A Meta-Review of Prevalence Meta-Analyses. Frontiers in Psychology, 12, 703838. 10.3389/fpsyg.2021.703838

23. Dölen, G., Darvishzadeh, A., Huang, K. W., & Malenka, R. C. (2013). Social reward requires coordinated activity of nucleus accumbens oxytocin and serotonin. Nature, 501(7466), 179–184. 10.1038/nature12518

24. Eberle, C., Fasig, T., Brüseke, F., & Stichling, S. (2021). Impact of maternal prenatal stress by glucocorticoids on metabolic and cardiovascular outcomes in their offspring: A systematic scoping review. PLOS ONE, 16(1), e0245386. 10.1371/journal.pone.0245386

25. Ellis, B. J., & Del Giudice, M. (2019). Developmental Adaptation to Stress: An Evolutionary Perspective. Annual Review of Psychology, 70(1), 111–139. 10.1146/annurev-psych-122216-011732

26. Fall, C. H. D. (2013). Fetal Programming and the Risk of Noncommunicable Disease. Indian Journal of Pediatrics, 80(Suppl. 1), S13–S20. 10.1007/s12098-012-0834-5

27. Faraji, J., & Metz, G. A. S. (2025). Harnessing BDNF Signaling to Promote Resilience in Aging. Aging and Disease, 16(4), 1813–1841. 10.14336/ad.2024.0961

28. Feng, X., Jia, M., Cai, M., Zhu, T., Hashimoto, K., & Yang, J. J. (2025). Central-peripheral neuroimmune dynamics in psychological stress and depression: insights from current research. Molecular Psychiatry, 30(10), 4881–4898. 10.1038/s41380-025-03085-y

29. Flannery, J. C., Tirrell, P. S., Baumgartner, N. E., & Daniel, J. M. (2025). Neuroestrogens, the hippocampus, and female cognitive aging. Hormones and Behavior, 170, 105710. 10.1016/j.yhbeh.2025.105710

30. Fowden, A. L., Valenzuela, O. A., Vaughan, O. R., Jellyman, J. K., & Forhead, A. J. (2016). Glucocorticoid programming of intrauterine development. Domestic Animal Endocrinology, 56, S121–S132. 10.1016/j.domaniend.2016.02.014

31. Frick, K. M., & Kim, J. (2018). Mechanisms underlying the rapid effects of estradiol and progesterone on hippocampal memory consolidation in female rodents. Hormones and Behavior, 104, 100–110. 10.1016/j.yhbeh.2018.04.013

32. Gallup (2021). Gallup Global Emotions 2021. Gallup. https://www.gallup.com/file/analytics/352337/Gallup_Global_Emotions_2021_Report.pdf

33. Georgousopoulou, V., Serdari, A., Manomenidis, G., & Dacou-Voutetakis, C. (2026). Prenatal stress, hormones, and fetal brain development: gender differences. Hormones, 25(1), 5–10. 10.1007/s42000-025-00696-6

34. Glynn, L. M., & Baram, T. Z. (2019). The influence of unpredictable, fragmented parental signals on the developing brain. Frontiers in Neuroendocrinology, 53, 100736. 10.1016/j.yfrne.2019.01.002

35. Glynn, L. M., Davis, E. P., Luby, J. L., Baram, T. Z., & Sandman, C. A. (2021). A predictable home environment may protect child mental health during the COVID-19 pandemic. Neurobiology of Stress, 14, 100291. 10.1016/j.ynstr.2020.100291

36. Gotlieb, N., Wilsterman, K., Finn, S. L., Browne, M. F., Bever, S. R., Iwakoshi-Ukena, E., et al. (2022). Impact of Chronic Prenatal Stress on Maternal Neuroendocrine Function and Embryo and Placenta Development During Early-to-Mid-Pregnancy in Mice. Frontiers in Physiology, 13, 886298. 10.3389/fphys.2022.886298

37. Grillo, C. A., Woodruff, J. L., Macht, V. A., & Reagan, L. P. (2019). Insulin resistance and hippocampal dysfunction: Disentangling peripheral and brain causes from consequences. Experimental Neurology, 318, 71–77. 10.1016/j.expneurol.2019.04.012

38. Gu, H., Lei, Y., Yao, Y., Chen, C., & Liu, C. (2025). Physiological and psychological responses to acute stress: A meta-analysis of the 171 studies of Trier Social Stress Test including 8452 healthy adults. Psychoneuroendocrinology, 180, 107566. 10.1016/j.psyneuen.2025.107566

39. Guan, S. z., Fu, Y. j., Zhao, F., Liu, H. y., Chen, X. h., Qi, F. q., et al. (2021). The mechanism of enriched environment repairing the learning and memory impairment in offspring of prenatal stress by regulating the expression of activity-regulated cytoskeletal-associated and insulin-like growth factor-2 in hippocampus. Environmental Health and Preventive Medicine, 26(1), 8. 10.1186/s12199-020-00929-7

40. Hales, C. N., & Barker, D. J. P. (1992). Type 2 (non-insulin-dependent) diabetes mellitus: the thrifty phenotype hypothesis. Diabetologia, 35(7), 595–601. 10.1007/bf00400248

41. Heck, A. L., & Handa, R. J. (2019). Sex differences in the hypothalamic–pituitary–adrenal axis’ response to stress: an important role for gonadal hormones. Neuropsychopharmacology, 44(1), 45–58. 10.1038/s41386-018-0167-9

42. Hernández-Vivanco, A., de la Vega-Ruiz, R., Montes-Mellado, A., Azcoitia, Í., & Méndez, P. (2025). Activational and Organizational Effects of Sex Hormones on Hippocampal Inhibitory Neurons. Journal of Neuroscience, 45(18), e1764242025. 10.1523/jneurosci.1764-24.2025

43. Holmes, A., & Wellman, C. L. (2009). Stress-induced prefrontal reorganization and executive dysfunction in rodents. Neuroscience & Biobehavioral Reviews, 33(6), 773–783. 10.1016/j.neubiorev.2008.11.005

44. Howland, M. A., Sandman, C. A., Davis, E. P., Stern, H. S., Phelan, M., Baram, T. Z., et al. (2021). Prenatal maternal mood entropy is associated with child neurodevelopment. Emotion, 21(3), 489–498. 10.1037/emo0000726

45. Hu, R. K., Zuo, Y., Ly, T., Wang, J., Meera, P., Wu, Y. E., et al. (2021). An amygdala-to-hypothalamus circuit for social reward. Nature Neuroscience, 24(6), 831–842. 10.1038/s41593-021-00828-2

46. Hurlemann, R., & Grinevich, V. (Eds.) (2018). Behavioral pharmacology of neuropeptides: Oxytocin. Springer. 10.1007/978-3-319-63739-6

47. Islam, M., Strawn, M., & Behura, S. K. (2022). Fetal origin of sex-bias brain aging. The FASEB Journal, 36(8), e22463. 10.1096/fj.202200255rr

48. Jagtap, A., Jagtap, B., Jagtap, R., Lamture, Y., & Gomase, K. (2023). Effects of Prenatal Stress on Behavior, Cognition, and Psychopathology: A Comprehensive Review. Cureus, 15(10), e47044. 10.7759/cureus.47044

49. Jensen Peña, C., Monk, C., & Champagne, F. A. (2012). Epigenetic Effects of Prenatal Stress on 11β-Hydroxysteroid Dehydrogenase-2 in the Placenta and Fetal Brain. PLoS ONE, 7(6), e39791. 10.1371/journal.pone.0039791

50. Kim, E. J., & Kim, J. J. (2023). Neurocognitive effects of stress: a metaparadigm perspective. Molecular Psychiatry, 28(7), 2750–2763. 10.1038/s41380-023-01986-4

51. Kirkpatrick, M., Mandal, G., Elhadidy, I., Mariani, N., Priestley, K., Pariante, C. M., et al. (2025). From placenta to the foetus: a systematic review of in vitro models of stress- and inflammation-induced depression in pregnancy. Molecular Psychiatry, 30(4), 1689–1707. 10.1038/s41380-024-02866-1

52. Lautarescu, A., Craig, M. C., & Glover, V. (2020). Prenatal stress: Effects on fetal and child brain development. International Review of Neurobiology, 151, 17–40. 10.1016/bs.irn.2019.11.002

53. Lazic, S. E. (2010). The problem of pseudoreplication in neuroscientific studies: is it affecting your analysis?. BMC Neuroscience, 11(1), 5. 10.1186/1471-2202-11-5

54. Lei, A. A., Phang, V. W. X., Lee, Y. Z., Kow, A. S. F., Tham, C. L., Ho, Y. C., et al. (2025). Chronic Stress-Associated Depressive Disorders: The Impact of HPA Axis Dysregulation and Neuroinflammation on the Hippocampus—A Mini Review. International Journal of Molecular Sciences, 26(7), 2940. 10.3390/ijms26072940

55. Lephart, E. D., Weber, K. S., & Hedges, D. W. (2026). Assessing the Roles of Aging, Estrogen, Nutrition, and Neuroinflammation in Women and Their Involvement in Alzheimer’s Disease—A Narrative Overview. International Journal of Molecular Sciences, 27(3), 1239. 10.3390/ijms27031239

56. Ma, Y., Liu, M., Yang, L., Zhang, L., Guo, H., Qin, P., et al. (2020). Loss of Estrogen Efficacy Against Hippocampus Damage in Long-Term OVX Mice Is Related to the Reduction of Hippocampus Local Estrogen Production and Estrogen Receptor Degradation. Molecular Neurobiology, 57(8), 3540–3551. 10.1007/s12035-020-01960-z

57. Markov, D. D. (2022). Sucrose Preference Test as a Measure of Anhedonic Behavior in a Chronic Unpredictable Mild Stress Model of Depression: Outstanding Issues. Brain Sciences, 12(10), 1287. 10.3390/brainsci12101287

58. Matyjek, M., Meliss, S., Dziobek, I., & Murayama, K. (2020). A Multidimensional View on Social and Non-Social Rewards. Frontiers in Psychiatry, 11, 818. 10.3389/fpsyt.2020.00818

59. McEwen, B. S., & Akil, H. (2020). Revisiting the Stress Concept: Implications for Affective Disorders. Journal of Neuroscience, 40(1), 12–21. 10.1523/jneurosci.0733-19.2019

60. McEwen, B. S., & Gianaros, P. J. (2011). Stress- and Allostasis-Induced Brain Plasticity. Annual Review of Medicine, 62(1), 431–445. 10.1146/annurev-med-052209-100430

61. McEwen, B. S., & Morrison, J. H. (2013). The Brain on Stress: Vulnerability and Plasticity of the Prefrontal Cortex over the Life Course. Neuron, 79(1), 16–29. 10.1016/j.neuron.2013.06.028

62. Mirzakhani, H. (2024). From womb to wellness: early environmental exposures, cord blood DNA methylation and disease origins. Epigenomics, 16(17), 1175–1183. 10.1080/17501911.2024.2390823

63. Mohebi, A., Pettibone, J. R., Hamid, A. A., Wong, J. M. T., Vinson, L. T., Patriarchi, T., et al. (2019). Dissociable dopamine dynamics for learning and motivation. Nature, 570(7759), 65–70. 10.1038/s41586-019-1235-y

64. Molet, J., Heins, K., Zhuo, X., Mei, Y. T., Regev, L., Baram, T. Z., et al. (2016). Fragmentation and high entropy of neonatal experience predict adolescent emotional outcome. Translational Psychiatry, 6(1), e702. 10.1038/tp.2015.200

65. Mori, F. K., & Shimosawa, T. (2025). The Fetal Environment and the Development of Hypertension— The Epigenetic Modification by Glucocorticoids. International Journal of Molecular Sciences, 26(1), 420. 10.3390/ijms26010420

66. Nyberg, L., Lövdén, M., Riklund, K., Lindenberger, U., & Bäckman, L. (2012). Memory aging and brain maintenance. Trends in Cognitive Sciences, 16(5), 292–305. 10.1016/j.tics.2012.04.005

67. O’Donnell, K. J., & Meaney, M. J. (2017). Fetal Origins of Mental Health: The Developmental Origins of Health and Disease Hypothesis. American Journal of Psychiatry, 174(4), 319–328. 10.1176/appi.ajp.2016.16020138

68. Olave, F. A., Aguayo, F. I., Román-Albasini, L., Corrales, W. A., Silva, J. P., González, P. I., et al. (2022). Chronic restraint stress produces sex-specific behavioral and molecular outcomes in the dorsal and ventral rat hippocampus. Neurobiology of Stress, 17, 100440. 10.1016/j.ynstr.2022.100440

69. Piao, X., Xie, J., & Managi, S. (2024). Continuous worsening of population emotional stress globally: universality and variations. BMC Public Health, 24(1), 3576. 10.1186/s12889-024-20961-4

70. Rodríguez Peris, L., Scheuber, M. I., Shan, H., Braun, M., & Schwab, M. E. (2024). Barnes maze test for spatial memory: A new, sensitive scoring system for mouse search strategies. Behavioural Brain Research, 458, 114730. 10.1016/j.bbr.2023.114730

71. Roth, T. L., Lubin, F. D., Funk, A. J., & Sweatt, J. D. (2009). Lasting Epigenetic Influence of Early-Life Adversity on the BDNF Gene. Biological Psychiatry, 65(9), 760–769. 10.1016/j.biopsych.2008.11.028

72. Russo, S. J., & Nestler, E. J. (2013). The brain reward circuitry in mood disorders. Nature Reviews Neuroscience, 14(9), 609–625. 10.1038/nrn3381

73. Serita, T., Fukushima, H., & Kida, S. (2017). Constitutive activation of CREB in mice enhances temporal association learning and increases hippocampal CA1 neuronal spine density and complexity. Scientific Reports, 7(1), 42528. 10.1038/srep42528

74. Slanzi, C. M., & Fernand, J. K. (2024). On the Use and Benefits of Electronic Data Collection Systems: A Tutorial on Countee. Behavior Analysis in Practice, 17(4), 1228–1237. 10.1007/s40617-024-00964-8

75. Spellman, T., Rigotti, M., Ahmari, S. E., Fusi, S., Gogos, J. A., & Gordon, J. A. (2015). Hippocampal– prefrontal input supports spatial encoding in working memory. Nature, 522(7556), 309–314. 10.1038/nature14445

76. Stevenson, K., Lillycrop, K. A., & Silver, M. J. (2020). Fetal programming and epigenetics. Current Opinion in Endocrine and Metabolic Research, 13, 1–6. 10.1016/j.coemr.2020.07.005

77. Sulyok, E., Farkas, B., & Bodis, J. (2023). Pathomechanisms of Prenatally Programmed Adult Diseases. Antioxidants, 12(7), 1354. 10.3390/antiox12071354

78. Szyf, M. (2013). DNA Methylation, Behavior and Early Life Adversity. Journal of Genetics and Genomics, 40(7), 331–338. 10.1016/j.jgg.2013.06.004

79. Tottenham, N. (2020). Early Adversity and the Neotenous Human Brain. Biological Psychiatry, 87(4), 350–358. 10.1016/j.biopsych.2019.06.018

80. Toufexis, D., Rivarola, M. A., Lara, H., & Viau, V. (2014). Stress and the Reproductive Axis. Journal of Neuroendocrinology, 26(9), 573–586. 10.1111/jne.12179

81. Walf, A. A., & Frye, C. A. (2007). The use of the elevated plus maze as an assay of anxiety-related behavior in rodents. Nature Protocols, 2(2), 322–328. 10.1038/nprot.2007.44

82. Wang, S., Shi, Y., Xin, R., Kang, H., Xiong, H., & Ren, J. (2025). Exploring the role of insulin resistance in bridging the metabolic syndrome and Alzheimer’s disease-a review of mechanistic studies. Frontiers in Endocrinology, 16, 1614006. 10.3389/fendo.2025.1614006

83. Wang, Z., & Storm, D. R. (2011). Maternal Behavior is Impaired in Female Mice Lacking Type 3 Adenylyl Cyclase. Neuropsychopharmacology, 36(4), 772–781. 10.1038/npp.2010.211

84. Weaver, I. C. G., Cervoni, N., Champagne, F. A., D’Alessio, A. C., Sharma, S., Seckl, J. R., et al. (2004). Epigenetic programming by maternal behavior. Nature Neuroscience, 7(8), 847–854. 10.1038/nn1276

85. Weinstock, M. (2017). Prenatal stressors in rodents: Effects on behavior. Neurobiology of Stress, 6, 3–13. 10.1016/j.ynstr.2016.08.004

86. Wong, S. D., Wright, K. P., Spencer, R. L., Vetter, C., Hicks, L. M., Jenni, O. G., et al. (2022). Development of the circadian system in early life: maternal and environmental factors. Journal of Physiological Anthropology, 41(1), 22. 10.1186/s40101-022-00294-0

87. Wu, T., Jia, X., Shi, H., Niu, J., Yin, X., Xie, J., et al. (2021). Prevalence of mental health problems during the COVID-19 pandemic: A systematic review and meta-analysis. Journal of Affective Disorders, 281, 91–98. 10.1016/j.jad.2020.11.117

88. Wu, X., Lu, C., Deng, Z., Xiao, W., Ni, H., & Zhao, C. (2025). Glucocorticoid exposure-induced alterations in epigenetic age from human preterm infants and human lung fibroblasts and hippocampal neuronal cells. Clinical Epigenetics, 17(1), 29. 10.1186/s13148-025-01837-9

89. Yagi, S., & Galea, L. A. M. (2019). Sex differences in hippocampal cognition and neurogenesis. Neuropsychopharmacology, 44(1), 200–213. 10.1038/s41386-018-0208-4

90. Yang, J., Jia, Y., Guo, T., Zhang, S., Huang, J., Lu, H., et al. (2025). Comparative Analysis of HPA-Axis Dysregulation and Dynamic Molecular Mechanisms in Acute Versus Chronic Social Defeat Stress. International Journal of Molecular Sciences, 26(13), 6063. 10.3390/ijms26136063

91. Yuan, K., Zheng, Y. B., Wang, Y. J., Sun, Y. K., Gong, Y. M., Huang, Y. T., et al. (2022). A systematic review and meta-analysis on prevalence of and risk factors associated with depression, anxiety and insomnia in infectious diseases, including COVID-19: a call to action. Molecular Psychiatry, 27(8), 3214–3222. 10.1038/s41380-022-01638-z

92. Zhou, S. B., Xue, M., Liu, W., Chen, Y. X., Chen, Q. Y., Lu, J. S., et al. (2023). Age-related attenuation of cortical synaptic tagging in the ACC is rescued by BDNF or a TrkB receptor agonist in both sex of mice. Molecular Brain, 16(1), 4. 10.1186/s13041-022-00992-x

93. Zohar, I., & Weinstock, M. (2011). Differential Effect of Prenatal Stress on the Expression of Cortiocotrophin-Releasing Hormone and its Receptors in the Hypothalamus and Amygdala in Male and Female Rats. Journal of Neuroendocrinology, 23(4), 320–328. 10.1111/j.1365-2826.2011.02117.x

94. Zoubovsky, S. P., Hoseus, S., Tumukuntala, S., Schulkin, J. O., Williams, M. T., Vorhees, C. V., et al. (2020). Chronic psychosocial stress during pregnancy affects maternal behavior and neuroendocrine function and modulates hypothalamic CRH and nuclear steroid receptor expression. Translational Psychiatry, 10(1), 6. 10.1038/s41398-020-0704-2

95. Zoubovsky, S. P., Williams, M. T., Hoseus, S., Tumukuntala, S., Riesenberg, A., Schulkin, J., et al. (2022). Neurobehavioral abnormalities following prenatal psychosocial stress are differentially modulated by maternal environment. Translational Psychiatry, 12(1), 22. 10.1038/s41398-022-01785-5

